# Brassinosteroids modulate autophagy through phosphorylation of RAPTOR1B by the GSK3-like kinase BIN2 in Arabidopsis

**DOI:** 10.1101/2022.02.21.481334

**Authors:** Ching-Yi Liao, Yunting Pu, Trevor M. Nolan, Christian Montes, Hongqing Guo, Justin W. Walley, Yanhai Yin, Diane C. Bassham

## Abstract

Macroautophagy/autophagy is a conserved recycling process that maintains cellular homeostasis during environmental stress. Autophagy is negatively regulated by TARGET OF RAPAMYCIN (TOR), a nutrient-regulated protein kinase that in plants is activated by several phytohormones, leading to increased growth. However, the detailed molecular mechanisms by which TOR integrates autophagy and hormone-signaling are poorly understood. Here, we show that TOR modulates brassinosteroid (BR)-regulated plant growth and stress-response pathways. Active TOR was required for full BR-induced growth in *Arabidopsis thaliana*. Autophagy was constitutively up-regulated upon blocking BR biosynthesis or signaling, and down-regulated by increasing the activity of the BR pathway. BRASSINOSTEROID-INSENSITIVE 2 (BIN2) kinase, a GSK3-like kinase functioning as a negative regulator in BR signaling, directly phosphorylated Regulatory-Associated Protein of TOR 1B (RAPTOR1B), a substrate-recruiting subunit in the TOR complex, at a conserved serine residue within a typical BIN2 phosphorylation motif. Mutation of RAPTOR1B serine 916 to alanine, to block phosphorylation by BIN2, repressed autophagy and increased phosphorylation of the TOR substrate autophagy-related protein 13a (ATG13a). By contrast, this mutation had only a limited effect on growth. We present a model in which RAPTOR1B is phosphorylated and inhibited by BIN2 when BRs are absent, activating the autophagy pathway. When BRs signal and inhibit BIN2, RAPTOR1B is thus less inhibited by BIN2 phosphorylation. This leads to increased TOR activity and ATG13a phosphorylation, and decreased autophagy activity. Our studies define a new mechanism by which coordination between BR and TOR signaling pathways helps to maintain the balance between plant growth and stress responses.

## Introduction

Organisms adapt to environmental changes by optimizing nutrient and energy usage. Macroautophagy/autophagy plays a role in this acclimation process by degradation and recycling of cellular components that are damaged or no longer needed, and in plants functions in stress responses, senescence, control of phytohormone signaling, and pathogen defense ^1, 2^. Autophagy is maintained at a low basal level in non-stressed conditions and is induced during senescence and environmental stress. Upon induction of autophagy, a cup-shaped vesicle called a phagophore expands to engulf the cargo that will be degraded and is sealed to generate a double-membrane autophagosome, which delivers the cargo into the vacuole for degradation and recycling ^2, 3^. Many conserved *AUTOPHAGY-RELATED* (*ATG*) genes encode proteins that function in autophagosome initiation and formation ^4^. Although initially thought to be a bulk degradation system, autophagy can be selective ^5^, in which specific receptors interact with ATG8 to recruit cargo to autophagosomes ^6, 7^. In plants, increasing evidence indicates that the interplay between autophagy (non-selective or selective) and hormone signaling pathways can assist in plant environmental adaptation ^1, 8–12^.

Studies in yeast and mammals have identified genes that function in the control of autophagy, although many of these do not exist in plants ^2^. A key negative regulator of autophagy is the TARGET OF RAPAMYCIN complex (TORC), a highly conserved Ser/Thr protein kinase complex that is present in yeast, animals, and plants ^13–16^. The TORC signaling pathway is critical for the coordination of cell division and growth, protein synthesis and metabolism, and repression of TORC is required for autophagy induction for nutrient remobilization and stress tolerance ^17, 18^. TORC in *Arabidopsis thaliana* contains the TOR catalytic subunit and two accessory subunits, LST8 (Lethal with Sec Thirteen 8), which stabilizes the TOR complex ^19^, and RAPTOR (Regulatory-Associated Protein), which recruits substrates and presents them to TOR for phosphorylation ^20, 21^. In Arabidopsis, two RAPTOR isoforms (RAPTOR1A and RAPTOR1B) have been identified and RAPTOR1B is the predominantly expressed one ^21, 22^. Disruption of TOR signaling leads to plant growth defects. Complete knockout of *TOR* in Arabidopsis is embryo-lethal, while downregulation of *TOR* arrests plant growth and induces autophagy ^14, 23, 24^. Knockout mutants in the TOR binding partners RAPTOR1B or LST8 also have growth defects and constitutive autophagy, which may be due to reduced efficiency of recruitment of TOR substrates and decreased protein complex stability, respectively ^22, 25–27^. Several TOR substrates have been identified in Arabidopsis, including the ribosomal p70 S6 kinase (S6K), which modulates growth through translational regulation ^28, 29^, and ATG13a, a component of the autophagy-initiating ATG1-ATG13 complex ^30, 31^.

TOR signaling modulates the activity of signaling pathways for phytohormones such as brassinosteroids (BRs) ^32, 33^. BRs are steroid hormones, which are essential for plant growth and development ^34^. Deficiency in BR biosynthesis or signaling leads to growth and developmental defects, including dwarfism due to disruption of cell elongation and cell division, de-etiolation in the dark (constitutive photomorphogenesis), and causes male sterility ^35–37^. The BR signaling pathway interacts with multiple other signaling pathways to regulate growth and stress responses (Nolan et al., 2020). BRs are perceived by the receptor BRASSINOSTEROID-INSENSITIVE 1 (BRI1) and the co-receptor BAK1 ^36, 38, 39^. The BR signal is transduced to downstream transcription factors through many signaling components ^40–42^; BRASSINOSTEROID-INSENSITIVE 2 (BIN2), a glycogen synthase kinase (GSK)-3-like kinase, acts as a critical negative component in the pathway ^43^. In the absence of BR, BIN2 phosphorylates the homologous transcription factors BRI1-EMS-Suppressor 1 (BES1) and BRASSINAZOLE-RESISTANT 1 (BZR1) ^44–47^. Phosphorylation by BIN2 inhibits BES1 and BZR1 functions through cytoplasmic retention, reduced DNA binding, and degradation via the proteasome and selective autophagy ^8, 33,48–50^. In the presence of BR, BES1 and BZR1 accumulate in the nucleus in an unphosphorylated form due to inhibition of BIN2 and likely the action of protein phosphatase 2A (PP2A) ^45, 48, 51^. Active BES1 and BZR1 then regulate the transcription of thousands of genes ^52–56^, leading to downstream BR responses.

We have previously demonstrated that BES1 is targeted for selective autophagy through interaction of its receptor DSK2 with ATG8, suggesting that selective autophagy functions in the regulation of growth and stress responses by BRs ^8^. It has also been shown that TOR inhibits degradation of phosphorylated BZR1 through suppression of autophagy ^33^, and BIN2 has been suggested to be a downstream effector of TOR in the regulation of growth ^32^. These findings suggest interaction between the BR, TOR signaling, and autophagy pathways. However, regulatory mechanisms involved in these complex interactions remain to be defined. Here we show that TOR can act downstream of BR signaling to modulate BR-induced plant growth and autophagy. Our results reveal that autophagy is inhibited when BR signaling is increased; and autophagy is activated when BR signaling is decreased. Moreover, we demonstrate that BIN2 phosphorylates RAPTOR1B at serine 916. A phosphorylation-null mutant of RAPTOR1B at this conserved serine residue increases ATG13a phosphorylation, leading to reduced autophagy activity without substantial effects on growth or S6K phosphorylation. BR signaling therefore regulates RAPTOR1B, potentially altering TOR substrate selection, to coordinate growth and stress responses in plants.

## Results

### TOR is required for BL-induced plant growth

Previous studies suggested that TOR regulates the BR signaling components BIN2 ^32^ and BZR1 ^33^. BRs regulate the expression of thousands of genes through transcription factors including BES1 and BZR1 ^57^, and TOR also functions in the regulation of gene expression in plants ^58^. To assess the extent to which the TOR and BR signaling pathways intersect via regulating the same set of genes, we compared the differentially expressed genes in the transcriptome of Arabidopsis WT (Col-0) plants after treatment with either brassinolide (BL), the most active BR, or AZD8055 (AZD), a TOR inhibitor, using previously published data generated by RNA-seq ^58, 59^. Clustering analysis and gene list comparisons demonstrated that 36% of genes downregulated after AZD treatment (conceptually TOR-upregulated) are upregulated by BL, while 29% of genes that were upregulated after AZD treatment (conceptually TOR-downregulated) showed downregulation upon BL treatment (Figure 1A and B). In contrast, only 7% of genes downregulated after AZD treatment were downregulated by BL and 14.6% genes upregulated after AZD treatment were induced by BL. The genes regulated by the TOR and BR signaling pathways have substantial overlap, and are regulated mostly in the same direction, which suggests that TOR and BR signaling interact in a cooperative manner.

**Figure 1.**
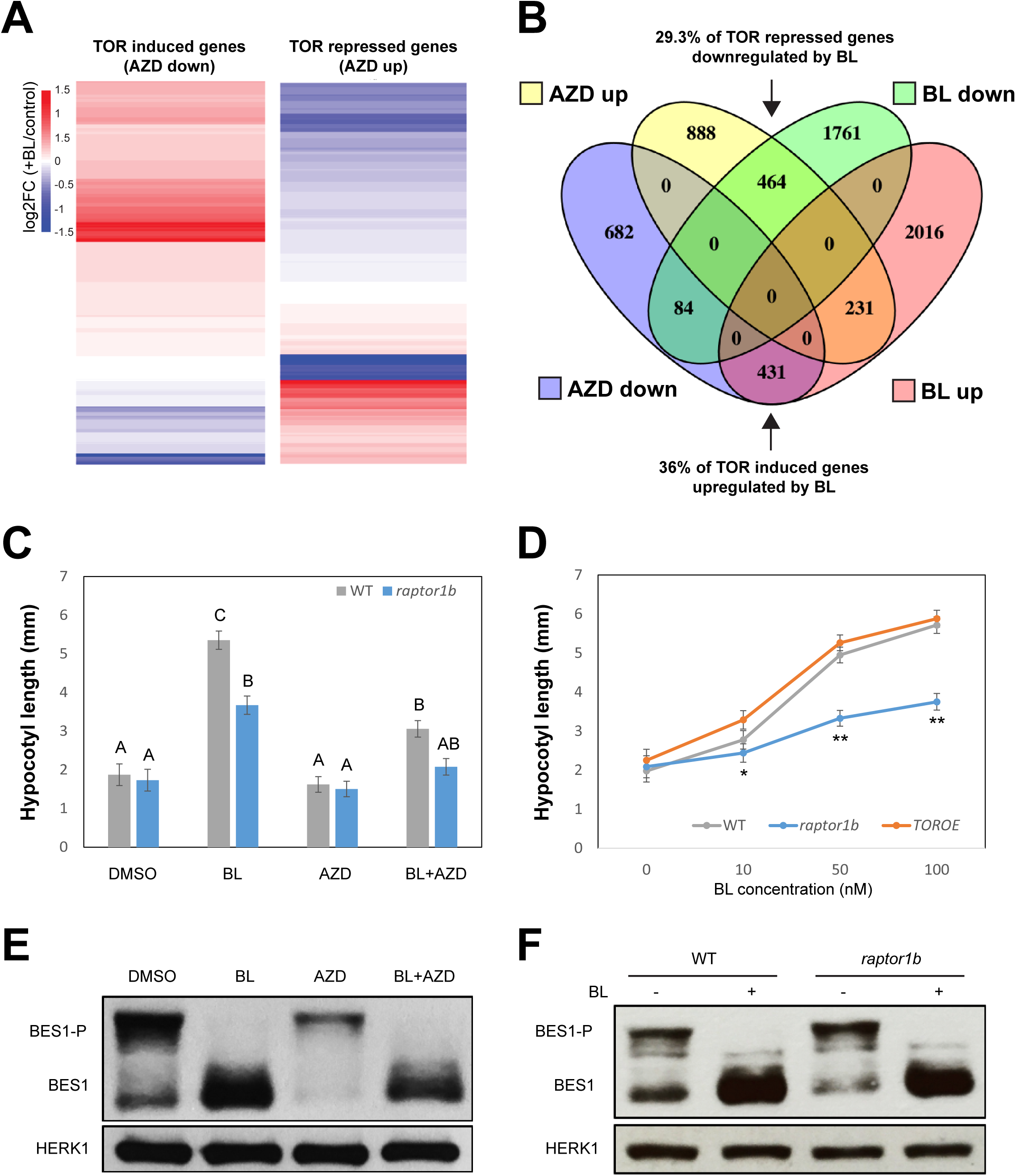
TOR is required for full BL-induced hypocotyl elongation. (**A**) Clustering analysis of TOR-regulated genes in response to BL treatment from whole transcriptome RNA-seq data. Previously reported RNA-seq data for 4-week-old WT plants treated with or without BL or AZD were used to construct heatmaps. BL, brassinolide; AZD, AZD8055. Color depicts log2 fold change upon BL treatment compared to the control. (**B**) Comparison of TOR- and BL-regulated genes in whole transcriptome RNA-seq data. Gene lists from (**A**) were compared using Venny. The AZD-downregulated transcriptome (AZD down) shows significant coregulation with the BL-upregulated transcriptome (BL up) (431/1197 genes, *p* -value: 9.49 × 10^-142^, Fisher’s exact test), while the AZD-upregulated transcriptome (AZD up) shows significant coregulation with the BL-downregulated transcriptome (BL down) (464/1583 genes, *p*-value: 3.00 × 10^-140^, Fisher’s exact test). (**C**) Hypocotyl elongation of WT and *raptor1b* seedlings in response to BL or AZD. Seedlings were grown on ½ MS medium with 100 nM BL, 1 μM AZD, or both chemicals for 7 days. (**D**) Hypocotyl elongation of WT, *raptor1b*, and *TOROE* seedlings in response to BL. Seedlings of each genotype were grown on ½ MS medium with the indicated BL concentration for 7 days. For panel (**C**) and (**D**), hypocotyl lengths of seedlings were measured and averaged from 15 independent seedlings and 3 independent replicates. Data represent means ± standard deviation (SD). In (**C**), different letters indicate statistically significant differences (*p* < 0.05) using Student’s *t*-test. In (**D**), asterisks indicate statistically significant differences (\*\**p* < 0.01, * *p* < 0.05) using Student’s *t*-test compared with WT under the same conditions. (**E**) Immunoblot of BES1 after BL or AZD treatment. WT seedlings were grown on solid ½ MS medium for 7 days and then transferred to solid ½ MS medium with DMSO or 1 μM AZD for 1 day, and transferred to liquid ½ MS medium with DMSO, 1 μM BL, 1 μM AZD, or both chemicals for 3 hours. (**F**) Immunoblot of BES1 upon BL treatment in WT and *raptor1b* mutant. WT and *raptor1b* seedlings were grown on solid ½ MS medium for 7 days and transferred to liquid ½ MS medium with DMSO or 1 μM BL for 2 hours. For panel (**E**) and (**F**), BES1 was detected via western blotting, with HERK1 as loading control. The experiments were repeated three times with similar results.

To determine if induction of plant growth by BR requires TOR, we analyzed the effect of the TOR inhibitor AZD on hypocotyl elongation in response to BL. Exogenous application of BL significantly increased hypocotyl length in WT seedlings, whereas AZD treatment alone had no obvious effect (Figure 1C). However, the addition of AZD reduced the elongation of hypocotyls in response to BL, indicating that inhibition of TOR signaling compromises BR-induced growth. To further test whether plants are less sensitive to BL upon disruption of TOR signaling, we assessed hypocotyl growth in response to BL of seedlings with genetic alterations of TOR activity. A TOR overexpression line (*TOROE*) ^13^, which has increased TOR activity, and a *raptor1b* knockout mutant ^21^, with decreased TOR activity, were exposed to BL and the effect on hypocotyl length measured and compared to WT seedlings. While the hypocotyl length of WT and *TOROE* seedlings increased with increasing concentration of BL, the BR response in *raptor1b* was significantly reduced at all tested concentrations of BL (Figure 1C and D).

In the absence of BR, BIN2 phosphorylates and inhibits BES1 functions, leading to inhibition of plant growth ^45^. We tested whether changes in BES1 phosphorylation cause the impaired BR response observed upon AZD treatment or in *raptor1b*. Immunoblotting showed that dephosphorylated BES1 accumulated upon BL treatment as expected, and co-treatment with AZD had little effect on BES1 phosphorylation (Figure 1E). Dephosphorylated BES1 also accumulated in both WT and *raptor1b* upon BL treatment (Figure 1F), indicating that blocking TOR signaling does not affect the BES1 phosphorylation pattern. We noticed that there is less total BES1 upon AZD treatment, possibly due to degradation by autophagy ^8^. Taken together, these results suggest that the BR and TOR signaling pathways may interact and have common outputs, that TOR can act downstream of BRs and is required for full BR-induced plant growth, and that TOR function in BR-regulated plant growth is largely independent of effects on BES1 phosphorylation.

### Autophagy is activated by BR inhibition and is suppressed when BR signaling is enhanced

Sugar starvation suppresses TOR activity and leads to autophagy induction in plants ^9, 60^, and combined with our observation that TOR is required for BR-induced growth, we hypothesized that BR might regulate autophagy. To test this hypothesis, we exposed 7-day-old seedlings expressing the autophagy marker GFP-ATG8e to BL or brassinazole (BRZ, a BR biosynthesis inhibitor) ^61^, and evaluated autophagosome numbers by counting GFP-labeled autophagosomes. Exogenous BL treatment significantly reduced the number of autophagosomes in plants upon sugar starvation compared to the control (Figure 2A), whereas inhibition of BR biosynthesis by BRZ increased the number of autophagosomes in control (+sucrose) conditions (Figure 2A). Vacuolar degradation of GFP-ATG8e after delivery by autophagy releases free GFP, and comparing the ratio of GFP-ATG8e to free GFP can therefore be used to measure flux through the entire autophagy pathway leading to degradation ^62^. Immunoblotting of BL- or BRZ-treated GFP-ATG8e-expressing plants using anti-GFP antibodies showed significant accumulation of free GFP in BRZ treated plants compared to the BL or control treatments, indicating higher autophagic flux upon BRZ treatment compared to the other conditions (Figure 2B). Upon sucrose starvation, the majority of GFP-ATG8e was processed to free GFP as expected.

**Figure 2.**
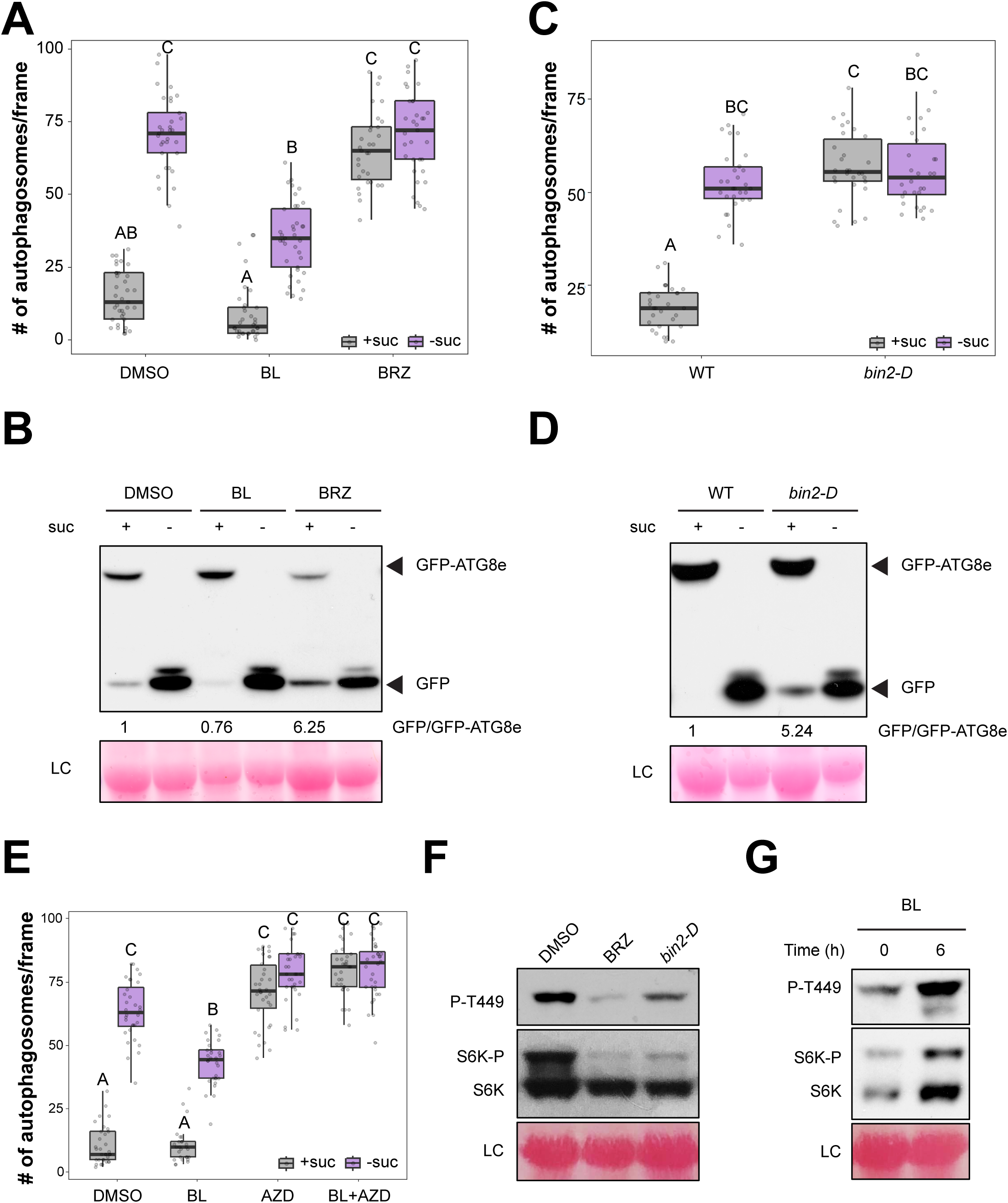
BL activates TOR and inhibits autophagy. (**A**) Number of GFP-tagged autophagosomes per frame in each condition. 7-day-old *GFP-ATG8e* seedlings were transferred to solid ½ MS medium under control (+suc) or starvation (-suc) conditions, plus 100 nM BL, 1 μM BRZ, or DMSO as control for an additional 3 days. (**B**) Immunoblot showing GFP-ATG8e cleavage using GFP antibody for plants as in (**A**). The ratio (the mean from all replicates) of GFP-ATG8e to free GFP is indicated with control condition set as 1 (-suc condition is not included because only free GFP was detected). Ponceau staining was used as a loading control (LC). (**C**) Number of GFP-tagged autophagosomes per frame in WT or *bin2-D* background. 7-day-old seedlings were transferred to solid ½ MS medium under control (+suc) or starvation (-suc) conditions for an additional 3 days. (**D**) Immunoblot showing GFP-ATG8e cleavage using GFP antibody for plants as in (**C**). The ratio (the mean from all replicates) of GFP-ATG8e to free GFP is indicated with control condition set as 1 (-suc condition is not included because only free GFP was detected). Ponceau staining was used as a loading control. (**E**) Number of GFP-tagged autophagosomes per frame in each condition. 7-day-old *GFP-ATG8e* expressing seedlings were transferred to solid ½ MS medium with control (+suc) or starvation (-suc), plus 100 nM BL, 1 μM AZD, both BL and AZD, or DMSO as control for an additional 3 days. (**F**) Immunoblot using S6K1/2 and P-T449-S6K antibodies of protein extracts from WT seedlings after treatment with 1 μM BRZ treatment for 7 days, and of 7-day-old *bin2-D* seedlings. (**G**) WT seedlings were grown on 1 μM BRZ for 7 days to suppress BL signaling, then transferred for 0 or 6 hours to 100 nM BL. Immunoblot was performed using S6K1/2 and P-T449-S6K antibodies after BL treatments. For (**A**), (**C**) and (**E**), GFP-labeled autophagosomes were observed by fluorescence microscopy and photographed. The average number of autophagosomes was calculated from at least 20 images per genotype for each condition. Data represent means ± SD. Boxes show the corresponding mean of each replicate with round dots representing individual measurements from each replicate. Different letters indicate statistically significant differences (*p* < 0.05) using Student’s *t*-test. Immunoblots were repeated a minimum of 3 times with similar results.

To further confirm that suppression of BR signaling leads to increased autophagy, we measured autophagosome numbers and GFP-ATG8e cleavage in *bin2-D* ^37^, a BIN2 gain-of-function mutant with reduced BR signaling, expressing GFP-ATG8e. Highly active autophagy was observed in control conditions when compared to WT plants (Figure 2C and D). We also assessed autophagy in additional BR mutants by transiently expressing *GFP-ATG8e* in protoplasts to label autophagosomes (Figure S1) ^63^. Under normal growth conditions, the percentage of protoplasts with active autophagy was increased in *bri1-301,* a partial loss-of-function BR receptor mutant, compared to WT. In contrast, upon sugar starvation the percentage of protoplasts with active autophagy was decreased in a BES1 gain-of-function mutant, *bes1-D*, which has enhanced BR signaling ^45, 54^. Our results indicate that suppression of BR signaling induces autophagy, and that conversely, activation of BR signaling reduces starvation-induced autophagy.

It is well established that TOR activation suppresses autophagy induction ^14–16^. Based on our observation that BR suppresses autophagy, we hypothesized that BR signaling activates the TOR signaling pathway, leading to inhibition of autophagy. Indeed, BL inhibited starvation-induced autophagy in *GFP-ATG8e* expressing seedlings (Figure 2E), whereas inhibition of TOR by AZD increased autophagy compared to the untreated conditions (Figure 2E). BL was unable to inhibit autophagy in the presence of AZD (Figure 2E), suggesting that BR signaling is upstream of TOR in the regulation of autophagy.

Next, we determined whether BR increases TOR kinase activity, thereby leading to suppression of autophagy. We investigated the phosphorylation status of the TOR substrate S6 protein kinase 1 (S6K1) in BRZ-treated seedlings and in *bin2-D* mutants. Blocking BR biosynthesis either with BRZ or in *bin2-D* reduced the amount of the S6K1 phosphorylated form (seen by a molecular weight shift on SDS-PAGE), and of Thr449-specific phosphorylation, a demonstrated *in vivo* target site of TOR kinase ^29^, seen using phospho-Thr449-specific S6K antibodies (Figure 2F). S6K1 phosphorylation was increased in WT seedlings by exogenous BL treatment (Figure 2G). Recently, Rodriguez et al. demonstrated that autophagy is rapidly induced within 30 minutes of various treatments, including addition of BL ^64^. Consistent with their findings, we observed that TOR activity rapidly decreased (Figure S2A) and autophagosome number increased (Figure S2B) for up to 1 hour of BL treatment. At later time points of 3 or 6 hours of treatment with BL, TOR activity increased and autophagy was repressed (Figure S2A and B). This indicates that the effect of BL on autophagy is dependent upon the time of exposure, and that after short-term activation of autophagy by BL to allow cell remodeling, BL reactivates TOR, leading to autophagy inhibition. These findings suggest that long-term treatment with BR represses autophagy through activation of TOR.

### Autophagy regulation by BR can be mediated by BIN2 through either TOR or BES1

Previous studies have shown that BIN2 phosphorylates a large number of substrates in addition to BES1 and BZR1 ^8, 65^. As the *bin2-D* mutant has an effect on both autophagy and TOR activity (Figure 2), we hypothesized that BIN2 acts upstream of TOR to control plant growth and autophagy. Since BIN2 also phosphorylates BES1 to regulate BL-related growth, we investigated whether BES1 is involved in affecting autophagy via BIN2. To test our hypothesis, we first transiently co-expressed mCherry-ATG8e as an autophagy marker with a YFP-BES1-D construct (BES1-dominant with P233L mutation) ^45^ in WT, *bin2-D,* or *raptor1b* protoplasts (Figure S3A and B), followed by sucrose starvation. As expected, in the non-starved control condition, protoplasts expressing mCherry-ATG8e only in a wild-type background had a basal level of autophagy, whereas in a *bin2-D* or *raptor1b* background, autophagy was increased. Expression of YFP-BES1-D had no effect on autophagy in any of the genotypes in control conditions. Upon starvation, expression of YFP-BES1-D in WT protoplasts suppressed the sugar starvation-induced autophagy, indicating that BES1 negatively regulates autophagy. However, expression of YFP-BES1-D in *bin2-D* or in *raptor1b* was unable to suppress their enhanced basal autophagy upon sugar starvation (Figure S3A and B). Similarly, we expressed YFP-BIN2-D in protoplasts from WT, *bes1-D*, or *TOROE* plants. Expression of YFP-BIN2-D in protoplasts from all three genotypes caused increased autophagy (Figure S3A and B). These findings suggest that BIN2-D constitutively activates autophagy even in the presence of enhanced BR signaling or in the TOR overexpression background, indicating that BIN2 is an autophagy activator and may regulate autophagy through multiple downstream pathways.

Based on the results from transient expression, we propose that BIN2 activates autophagy via two separate branches: one branch involves BIN2-TORC and the other branch involves BIN2-BES1. To reveal genetic relationships between *BIN2*, *BES1*, *TOR*, and *RAPTOR1B*, we generated *bes1-D bin2-D*, *bes1-D raptor1b,* and *TOROE bin2-D* double mutants from the corresponding single mutants with either increased or decreased BR or TOR signaling, and examined their autophagy and growth phenotypes (Figure 3A and B). We transiently expressed *GFP-ATG8e* as an autophagy marker in protoplasts from each genotype. Consistent with the transient expression results shown in Figure S3, the *bes1-D* mutant and *TOROE* protoplasts had reduced autophagy under sucrose starvation conditions, to a level comparable to WT under the non-starvation condition (Figure 3A). In contrast, *bin2-D* and *raptor1b* protoplasts had constitutive autophagy even in non-starvation conditions (Figure 3A). The *bes1-D bin2-D* double mutant had constitutive autophagy (Figure 3A), suggesting that, in the double mutant, BIN2-D still inhibits TORC, leading to up-regulation of autophagy even in the presence of constitutively active BES1. The constitutive autophagy observed in the *bes1-D raptor1B* double mutant supports the possibility that BIN2 regulates TORC independently of BES1. These double mutant phenotypes suggest that BIN2 regulates autophagy through TORC and BES1 independently. Notably, the *TOROE bin2D* double mutant displayed a constitutive autophagy phenotype (Figure 3A). The most likely explanation is that the substantially up-regulated BIN2 kinase activity in *bin2-D* can inhibit the slightly overexpressed TOR in the double mutant. It is also conceivable that in the *TOROE bin2-D* double mutant, BIN2 inhibits BES1, leading to constitutive autophagy through the BIN2-BES1 branch, independent of the TORC branch and overcoming any TOR overexpression effect on autophagy. These findings together support the conclusion that BIN2 modulates autophagy through multiple pathways.

**Figure 3.**
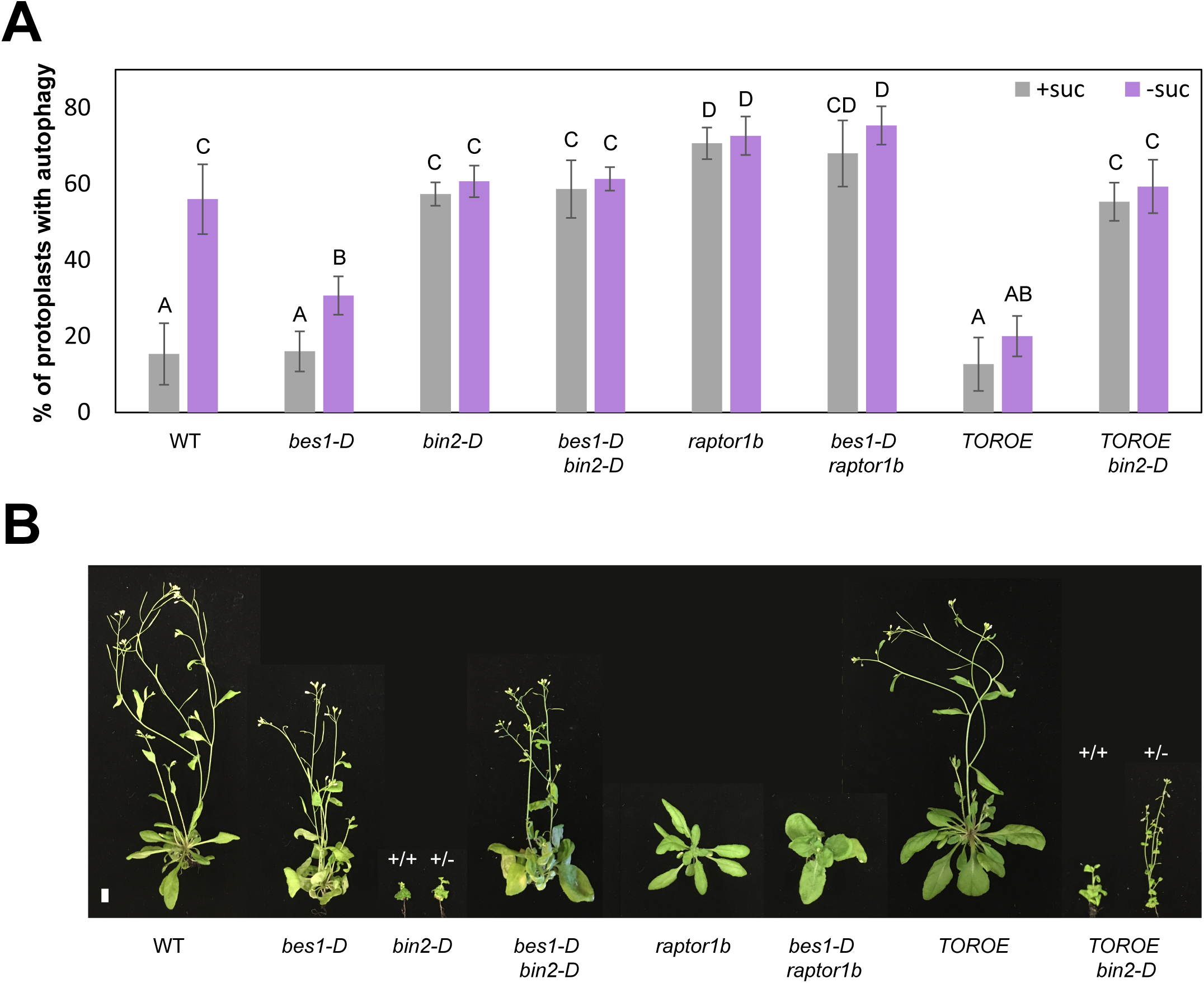
BIN2 activates autophagy and inhibits plant growth. (**A**) Quantification of autophagy in protoplasts by calculating the percentage of protoplasts with 3 or more GFP-ATG8e-labeled autophagosomes in the indicated genotype for each condition. Data represent means ± SD from 3 biological replicates. Different letters indicate statistically significant differences (*p* < 0.05) using Student’s *t*-test. (**B**) Phenotype of 40-day-old representative adult plants of each genotype. Scale bar = 10 mm.

Interestingly, the growth phenotypes of double mutants *TOROE bin2-D*, *bes1-D bin2-D*, and *bes1-D raptor1b* were either in between their parent genotypes or similar to one parent. *bes1-D* was dominant in showing curly leaves and early senescence in both *bes1-D bin2-D* and *bes1-D raptor1b* (Figure 3B), but the later flowering time observed in both double mutants was more like their *bin2-D* or *raptor1b* parent (Figure 3B). Meanwhile, *bin2-D* was dominant in showing most of the *bin2-D* growth phenotype in *TOROE bin2-D* (Figure 3B). These results suggest that BIN2 regulates autophagy and growth partially through different pathways, which is consistent with the overlap between BR and TOR-regulated genes (Figure 1A and B). Taken together, our findings suggest that BIN2 regulates autophagy by inhibiting either TOR or BES1.

### BIN2 phosphorylates and directly interacts with RAPTOR1B

Our results suggest that BIN2 kinase affects TOR activity, which in turn controls the activity of the autophagy pathway. BIN2 is a critical regulator of many aspects of plant growth, development, and stress responses via protein phosphorylation. We therefore hypothesized that BIN2 phosphorylates proteins in the TOR pathway, thus regulating TOR activity. To determine whether BIN2 phosphorylates TOR complex components, we analyzed a previously reported proteome-wide dataset of BIN2 direct targets generated using the *in vitro* multiplexed assay for kinase specificity (MAKS) ^66–68^. We also examined *in vivo* phosphoproteome data for sites differentially phosphorylation in *bin2-D* plants ^66^. In the MAKS assay, we identified the phosphopeptides TPPVpSPPR and TPPVpSPPRTNYLSGLR from RAPTOR1B, which contain the phosphosite Ser916, as significantly increased compared to the control samples with no BIN2 added (Figure 4A). In the *in vivo* phosphoproteome comparisons, Ser916 phosphorylation was significantly decreased in BIN2 loss-of-function *bin2-3 bil1 bil2* triple mutants (*bin2-T*) compared to *bin2-D*. We therefore propose that BIN2 phosphorylates RAPTOR1B at Ser916, leading to the autophagy activation we observed in *bin2-D*. Ser916 is located between the RAPTOR1B HEAT repeats and WD40 domain and is predicted to be exposed on the surface of the TOR complex (based on the crystal structure of the mammalian TOR complex) ^69^ (Figure 4B).

**Figure 4.**
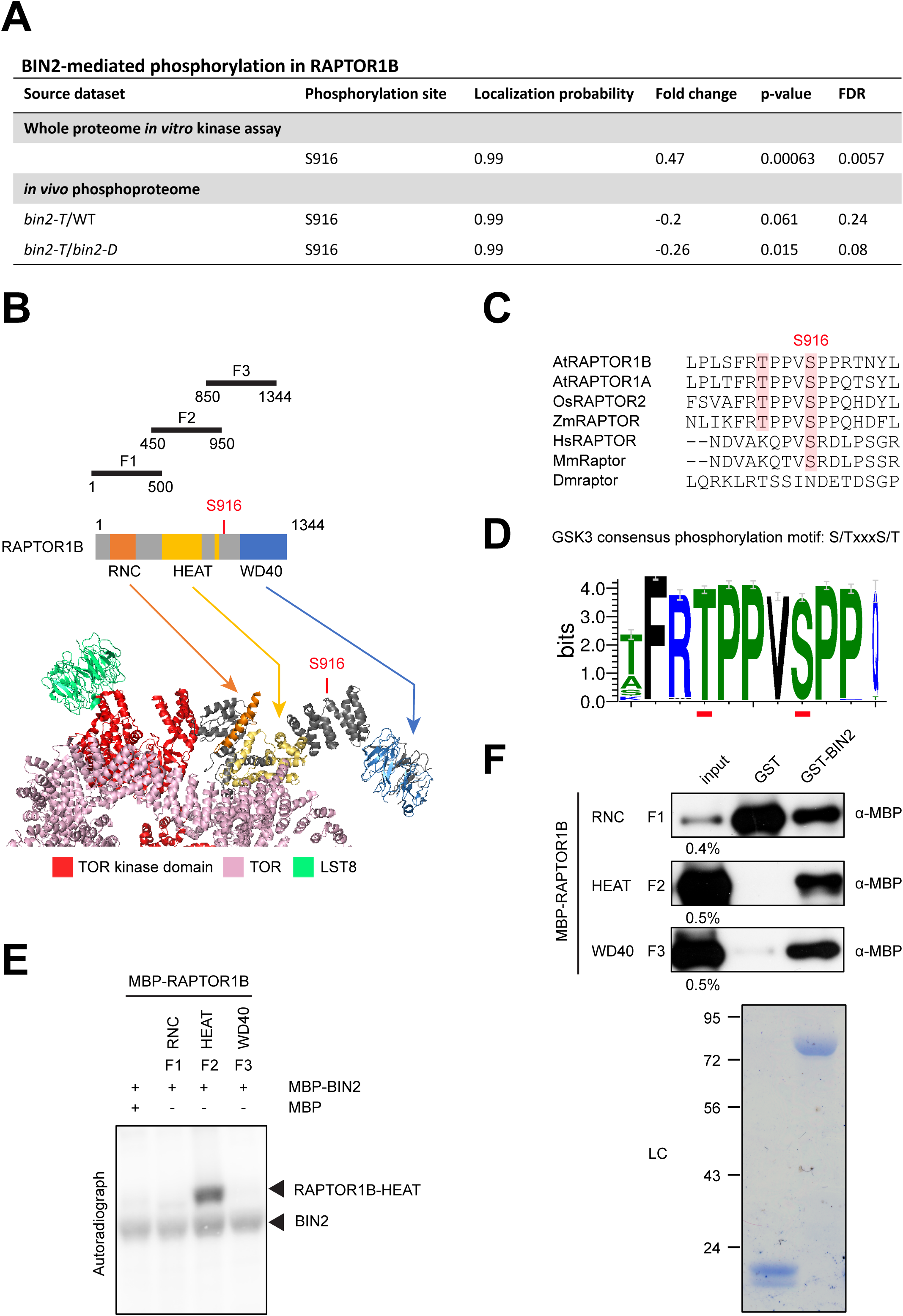
BIN2 phosphorylates and interacts with RAPTOR1B *in vitro*. (**A**) Detection of BIN2-mediated phosphorylation of RAPTOR1B from a whole proteome *in vitro* kinase assay and *in vivo* phosphoproteome (*bin2-T* compared to WT and *bin2-T* compared to *bin2-D*). Phosphopeptides which include serine 916 (S916) were significantly upregulated in phosphoproteome datasets. Fold change, log2 fold change. (**B**) Schematic representation of Arabidopsis RAPTOR1B. The position of the RNC domain is highlighted in orange, HEAT domain in yellow, and WD40 domain in blue. Fragments used to make recombinant MBP-RAPTOR1B proteins are shown, with the RNC domain-containing fragment named F1, HEAT domain F2, and WD40 domain F3. A partial Arabidopsis TOR complex 3D structure predicted based on mTOR was generated by PyMOL, color coded the same as the schematic. (**C**) Alignment of RAPTOR1B protein sequences surrounding the S916 phosphosite. Sequences from representative eukaryotic species are shown: *Arabidopsis thaliana* (At), *Drosophila melanogaster* (Dm), *Homo sapiens* (Hs), *Mus musculus* (Mm), *Oryza sativa* (Os), and *Zea mays* (Zm). (**D**) Weblogo created from multiple alignment of 250 closest-related plant RAPTOR1B paralogues shows the consensus sequence among species. (**E**) Phosphorylation of RAPTOR1B by BIN2 *in vitro*. Recombinant MBP-RAPTOR1B-RNC (F1), MBP-RAPTOR1B-HEAT (F2), and MBP-RAPTOR1B-WD40 (F3) proteins were purified from *E. coli* and incubated with MBP-BIN2 in kinase buffer containing γ-^32^P-ATP, then separated by SDS-PAGE. RAPTOR1B phosphorylation (^32^P-RAPTOR1B) was detected by autoradiography. (**F**) GST pull-down showing interactions of GST-BIN2 or GST as control with F2 and F3. MBP-RAPTOR1B fragments were detected with anti-MBP antibody. LC, loading control indicates amounts of MBP proteins used in pull-down reactions. Experiments were repeated a minimum of 3 times with similar results.

BIN2 is a member of the highly conserved GSK3-like kinase family, which recognizes the consensus sequence S/T-X-X-X-S/T. RAPTOR1B Ser916 is part of a GSK consensus motif (along with either Thr912 or Thr920). We analyzed RAPTOR protein sequences from several representative eukaryotes (Figure 4C) and also compared 250 of the most closely-related plant RAPTOR1B paralogues to assess whether these GSK motifs are conserved among species. We found that Ser916 is conserved in most plant species including *Arabidopsis thaliana*, *Oryza sativa*, and *Zea mays* (Figure 4C and D). Although Ser916 is also conserved in mammals such as *Homo sapiens* and *Mus musculus*, there are no GSK motifs surround the conserved serine found in these organisms, suggesting that BIN2/GSK3 phosphorylation of RAPTOR could be plant-specific (Figure 4C).

To further validate the direct phosphorylation of RAPTOR1B by BIN2, we generated recombinant proteins containing 3 different RAPTOR1B fragments (Figure 4B) and tested whether they can be phosphorylated by recombinant BIN2 *in vitro*. Phosphorylation by BIN2 was detected for RAPTOR1B-HEAT (F2) but not for RAPTOR1B-RNC (F1) or RAPTOR1B-WD40 (F3), indicating that BIN2 phosphorylates RAPTOR1B on the RAPTOR1B fragment containing the HEAT domain *in vitro* (Figure 4B and E). Because the majority of the RAPTOR1B-WD40 protein (F3) was degraded (Figure S4), we were unable to conclusively determine whether BIN2 phosphorylates F3. Given that BIN2 phosphorylates RAPTOR1B *in vitro*, we asked whether BIN2 and RAPTOR1B interact directly. We tested for a potential direct interaction between GST-BIN2 and the three RAPTOR1B recombinant protein fragments by glutathione S-transferase (GST) pull-down assays (Figure 4F). RAPTOR1B fragments containing HEAT (F2) or WD40 (F3) domains both interacted with GST-BIN2 (Figure 4F). Because the GST alone bound strongly to the RAPTOR1B RNC domain (F1), we were unable to determine whether GST-BIN2 also interacts with F1 (Figure 4F). Nevertheless, these results showed a direct interaction between BIN2 and RAPTOR1B, supporting the idea that BIN2 interacts with RAPTOR1B and regulates it through phosphorylation.

### Phosphorylation of RAPTOR1B serine 916 by BIN2 modulates autophagy

Our findings suggest that BIN2 directly interacts with and phosphorylates the RAPTOR1B subunit of the TOR complex. We hypothesized that the phosphorylation of Ser916 of RAPTOR1B by BIN2 activates autophagy due to the inhibition of TOR signaling, whereas loss of phosphorylation at this site should promote plant growth and repress autophagy. To test this hypothesis, we generated transgenic plants carrying either 35S promoter-driven wild-type *RAPTOR1B* or a non-phosphorylatable *RAPTOR1B^S916A^*mutant, both in a *raptor1b* mutant background. Transgenic plants that have similar expression levels were identified and grouped for comparison; no major difference in phenotype was seen between groups (Figure S5A). Disruption of *RAPTOR1B* broadly delays plant development in Arabidopsis, including slow germination and root growth, and delayed bolting ^27, 70^. In all 3 independent lines of *WT RAPTOR1B* or *RAPTOR1B^S916A^*with different expression levels, transgenic expression of wild-type or mutated *RAPTOR1B* complemented the *raptor1b* phenotypes in germination, root elongation, and plant growth under normal growth conditions, indicating that both WT RAPTOR1B and RAPTOR1B^S916A^ is functional in plant growth. No significant differences were observed between growth of *raptor1b* expressing the *WT RAPTOR1B* or *RAPTOR1B^S916A^*, or between their responses to BL (Figure S5B and C), indicating that BIN2 phosphorylation of RAPTOR1B on S916 does not affect growth under normal conditions. By contrast, the hypocotyls of dark-grown *RAPTOR1B^S916A^* in all groups were slightly longer than those of the wild-type control seedlings or *raptor1b* plants expressing wild-type *RAPTOR1B* (Figure S5D). Although *raptor1b* was less sensitive to BL and BRZ in the dark compared to wild-type seedlings (Figure S5C and D), we did not find a significant difference in hypocotyl length in response to BL or BRZ between wild-type or *raptor1b* plants expressing wild-type *RAPTOR1B* or *RAPTOR1B^S916A^*(Figure S5C and D). These findings suggest that the Ser916 may not contribute substantially to BR-regulated plant growth under normal conditions.

Since the RAPTOR1B subunit of TORC acts in recruiting substrates for TOR kinase, we investigated whether BIN2 phosphorylation of RAPTOR1B Ser916 modulates TOR activity. When activated, TOR phosphorylates S6K1/2 to promote plant growth. Immunoblotting showed that S6K phosphorylation was undetectable in *raptor1b*, and S6K protein level was also significantly reduced in *raptor1b* compared to WT (Figure S5E). We did not detect a significant difference in S6K-PT449 phosphorylation or S6K phosphorylation in *raptor1b* expressing *RAPTOR1B^S916A^* compared with either WT or *raptor1b* carrying wild-type *RAPTOR1B* (Figure S5E), consistent with the lack of a major effect on growth.

To assess whether phosphorylation of RAPTOR1B at Ser916 by BIN2 regulates autophagy, we evaluated how mutation of the phosphorylation site affects autophagy *in planta*. First, we transiently co-expressed *WT RAPTOR1B* or *RAPTOR1B^S916A^*, or a vector control with GFP-ATG8e as an autophagy marker in *raptor1b* protoplasts. Expression of *RAPTOR1B^S916A^* in protoplasts from *raptor1b* repressed autophagy (Figure S6A), indicating that *RAPTOR1B^S916A^* is functional in repressing the constitutive autophagy seen in *raptor1b*. We hypothesized that BIN2 phosphorylates RAPTOR1B at S916 to repress autophagy. To evaluate whether *RAPTOR1B^S916A^* is able to repress constitutive autophagy in *bin2-D* plants, we transiently co-expressed *GFP-ATG8e* with vector control, *WT RAPTOR1B,* or *RAPTOR1B^S916A^* in *bin2-D* protoplasts. Expression of the *WT RAPTOR1B* in protoplasts from *bin2-D* decreased the constitutive autophagy in *bin2-D*, while expression of *RAPTOR1B^S916A^*in protoplasts from *bin2-D* showed a much stronger repression than the effect of the *WT RAPTOR1B* (Figure S6A). Upon overexpression of *RAPTOR1B^S916A^*, most of the RAPTOR1B in *bin2-D* protoplasts would be the non-phosphorylatable form at Ser916, leading to a stronger repression of autophagy.

We generated transgenic plants co-expressing *GFP-ATG8e* and the *WT RAPTOR1B* or *RAPTOR1B^S916A^* in the *raptor1b* mutant background, in lines expressing “Moderate” levels of *RAPTOR1B*, similar to that in wild-type plants. Autophagosome number was measured in each genotype. These results showed that loss of Ser916 phosphorylation in RAPTOR1B partially suppressed the induction of autophagy upon blocking BR synthesis or sugar starvation compared to the control condition (Figure 5A). The *raptor1b* mutant had increased numbers of autophagosomes compared with WT seedlings, and *WT RAPTOR1B* complemented this phenotype of *raptor1b* and gave similar autophagy levels as those in WT plants in response to BL, BRZ or sugar starvation (Figure 5A). Transgenic expression of *RAPTOR1B^S916A^* complemented the enhanced autophagy of the *raptor1b* mutant (Figure 5A). Interestingly, *RAPTOR1B^S916A^* expression led to reduced autophagy compared with WT in response to BRZ or sucrose starvation (Figure 5A), suggesting that while blocking BR synthesis or starving for sucrose induces autophagy in seedling roots, these treatments have less effect in the non-phosphorylatable *RAPTOR1B^S916A^* transgenic plants.

**Figure 5.**
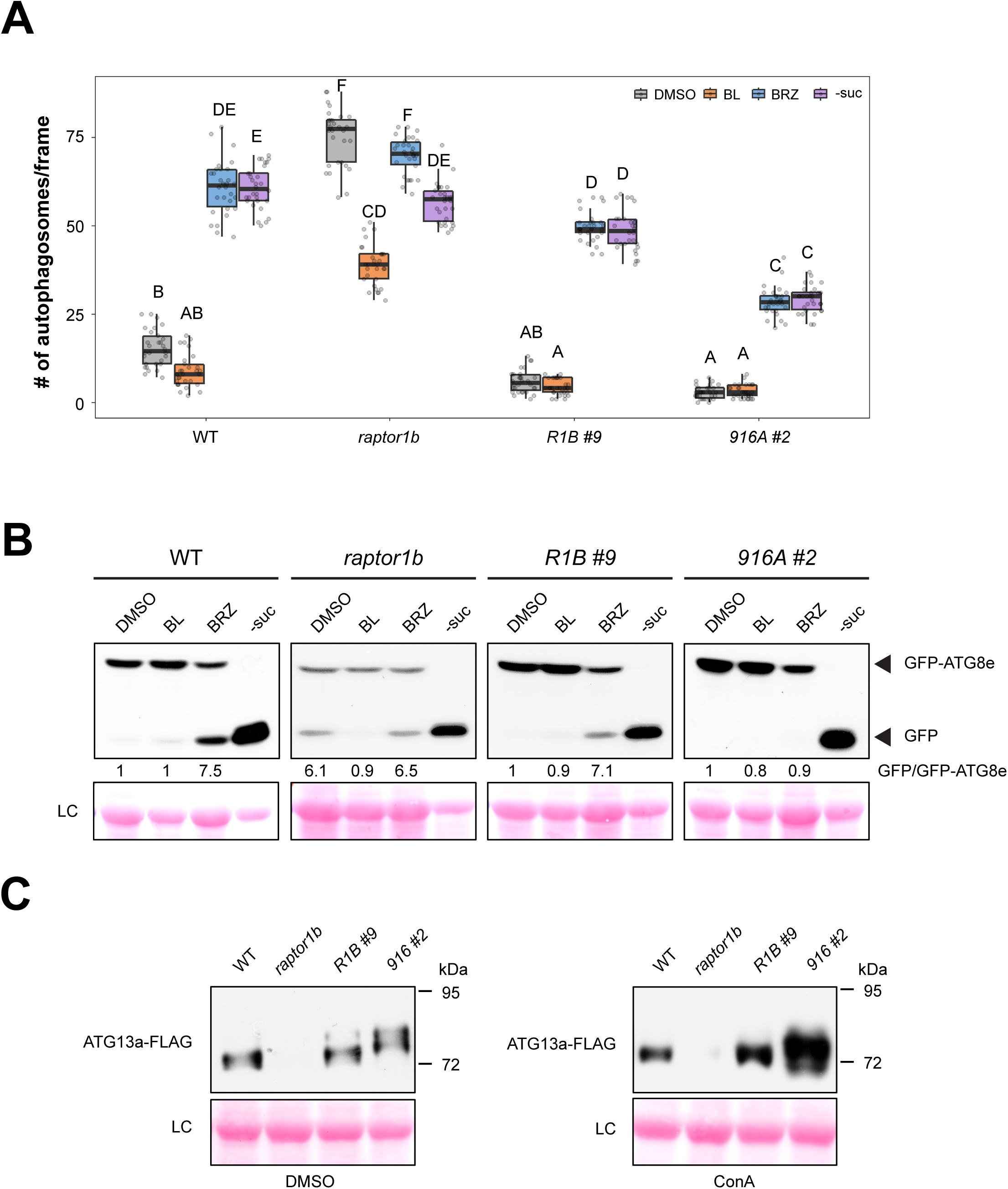
Phosphorylation of RAPTOR1B serine 916 by BIN2 modulates autophagy. (**A**) Number of GFP-tagged autophagosomes per frame for the indicated genotypes for each condition. 7-day-old GFP-ATG8e-expressing seedlings of each genotype were transferred to solid ½ MS medium under control or starvation (-suc) conditions, plus 100 nM BL, 1 μM BRZ, or DMSO as control for an additional 3 days. GFP-labeled autophagosomes were observed by fluorescence microscopy and photographed. The average number of autophagosomes was calculated from at least 20 images per genotype for each condition. Data represent means ± SD. Boxes show the corresponding mean of each replicate with round dots representing individual measurements from each replicate. Different letters indicate statistically significant differences (*p* < 0.05) using Student’s *t*-test. (**B**) Immunoblot showing GFP-ATG8e cleavage using GFP antibody for plants as in (**A**). The ratio (the mean from all replicates) of GFP-ATG8e to free GFP is indicated with control condition set as 1 (-suc condition is not included because only free GFP was detected). Ponceau staining was used as a loading control. (**C**) Immunoblot of ATG13a-FLAG phosphorylation in WT, *raptor1b*, and transgenic plants expressing *WT RAPTOR1B* (*R1B #9*) or *RAPTOR1B^S916A^*(*916A #2*) in a *raptor1b* background. Leaf protoplasts prepared from the indicated genotypes, transiently expressing ATG13a-FLAG, were treated with DMSO or 1 μM concanamycin A (ConA) for 12 hours. Ponceau staining was used as a loading control. Experiments were repeated a minimum of 3 times with similar results.

We also counted the number of fluorescent puncta in 3 independent lines of *WT RAPTOR1B* or *RAPTOR1B^S916A^*after staining with monodansylcadaverine (MDC), which labels acidic vesicles including autophagosomes. These results were consistent with the GFP-ATG8e quantification and no major differences between lines with different RAPTOR1B or *RAPTOR1B^S916A^* expression levels were seen (Figure S6B).

Immunoblotting of GFP-ATG8e-expressing plants treated as in Figure 5A using anti-GFP antibodies showed significant accumulation of free GFP in BRZ treated plants and *raptor1b* compared to the BL or control treatments, indicating higher autophagic flux upon BRZ treatment and in *raptor1b* compared to the other conditions (Figure 5B). Accumulation of free GFP was significantly reduced in BL treated *raptor1b* and BRZ treated *RAPTOR1B^S916A^* (Figure 5B), which was consistent with our microscopic observations. These data demonstrate that enhanced BL signaling inactivates BIN2, which causes loss of phosphorylation at RAPTOR1B Ser916 and suppresses autophagy.

While TOR phosphorylates S6K1/2 to promote plant growth, a major substrate in inhibition of autophagy is the autophagy initiation complex component ATG13. To determine whether phosphorylation of RAPTOR1B by BIN2 affected TOR activity towards ATG13, we transiently expressed *ATG13a-FLAG* in protoplasts from each genotype. Strikingly, the higher molecular mass (> 80-kD) ATG13a-FLAG phosphorylated forms were significantly increased in *raptor1b* expressing *RAPTOR1B^S916A^* compared with either WT or *raptor1b* carrying *WT RAPTOR1B* (Figure 5C). ATG13a can be degraded by autophagy under nutrient-rich conditions ^31^. Addition of the V-ATPase inhibitor concanamycin A (ConA), which blocks autophagy-mediate degradation, increased the amount of detectable ATG13a-FLAG protein and ATG13a-FLAG phosphorylated forms in *raptor1b* carrying *RAPTOR1B^S916A^*compared with other controls (Figure 5C). Overall, these findings suggest that loss of phosphorylation at RAPTOR1B Ser916 leads to an increase in ATG13a phosphorylation level, but not in that of S6K, thereby inactivating autophagy but having a minimal effect on growth.

## Discussion

Plants have evolved complex signaling networks to coordinate anabolism and catabolism for survival under constantly changing environmental conditions. TOR plays a central role in this coordination by activating growth and repressing stress responses ^71^. BRs are a family of plant hormones that also stimulate plant development and interact with multiple signaling pathways including those involved in stress tolerance ^34, 72^. We previously showed that under stress conditions, the BR signaling component BES1 is targeted for selective autophagy through the ubiquitin receptor DSK2 ^8^. Here we show that the protein kinase BIN2 acts upstream of the TOR complex and inhibits its activity, thereby promoting autophagy. Our findings further reveal that BR signaling regulates autophagy via BIN2 phosphorylation of RAPTOR1B, the substrate recruiting subunit in the TOR complex, affecting the downstream signaling that controls nutrient recycling in plants.

The GLYCOGEN SYNTHASE KINASE3 (GSK3) family is a highly conserved serine/threonine kinase family which is involved in a broad range of biological processes in eukaryotes ^73^. BIN2 is one of the best-studied GSK3-like kinases in Arabidopsis ^74^. In mammals, phosphorylation of RAPTOR by GSK3 at Ser859 increases mTOR activity and triggers mTOR-mediated amino acid-dependent signaling ^75^. However, direct regulation of the TOR complex via GSK3 in plants has not previously been shown ^74^. Typically, phosphorylation events mediated by GSK3 occur in the consensus sequence S/T-X-X-X-S/T. This sequence is present around Ser916 of RAPTOR1B in Arabidopsis and is conserved across RAPTOR1B orthologs in plants (Figure 4C). Furthermore, the location of Ser916 on the surface of the TOR complex makes its phosphorylation by BIN2 feasible (Figure 4B). BIN2 may regulate TOR substrate recruitment efficiency or substrate selectivity through direct phosphorylation at the RAPTOR1B Ser916 site. Analysis of a recently published BL time-series dataset ^47^ indicated that Ser916 phosphorylation status varies in response to different BL treatment times. Consistent with our observations in Figure S2, Ser916 phosphorylation rapidly increased in the beginning of BL treatment and decreased at the later time points. Interestingly, unlike Ser916, a change in phosphorylation of the plant-conserved Thr912 residue was only detected in some of the tested BL treatment time points. Ser916 phosphorylation level therefore is correlated with regulation of autophagy by TOR in response to BR.

BR signaling may play an important role in coordinating TOR signaling and autophagy in response to environmental cues. We note that both phosphorylation and protein abundance of S6K, a well-known TOR substrate that controls translation and growth ^60^, are not substantially changed in a RAPTOR1B Ser916-to-Ala non-phosphorylatable mutant, which is consistent with the minor effect on growth. Meanwhile, the phosphorylation status of another TOR target, ATG13a, a component of the autophagy initiation complex, is substantially increased in a RAPTOR1B Ser916 phosphorylation null mutant, in which autophagy is repressed. These observations suggest that Ser916 phosphorylation by BIN2 may primarily affect autophagy but not growth. We propose a model (Figure 6) in which, in the presence of BR, BIN2 is inhibited, which leads to a decrease in RAPTOR1B Ser916 phosphorylation. Decreased Ser916 phosphorylation on RAPTOR1B promotes phosphorylation of ATG13a by TOR, thus repressing autophagy (Figure 6). When BR is absent, BIN2 can suppress the TOR complex through phosphorylation of RAPTOR1B and therefore induce autophagy.

**Figure 6.**
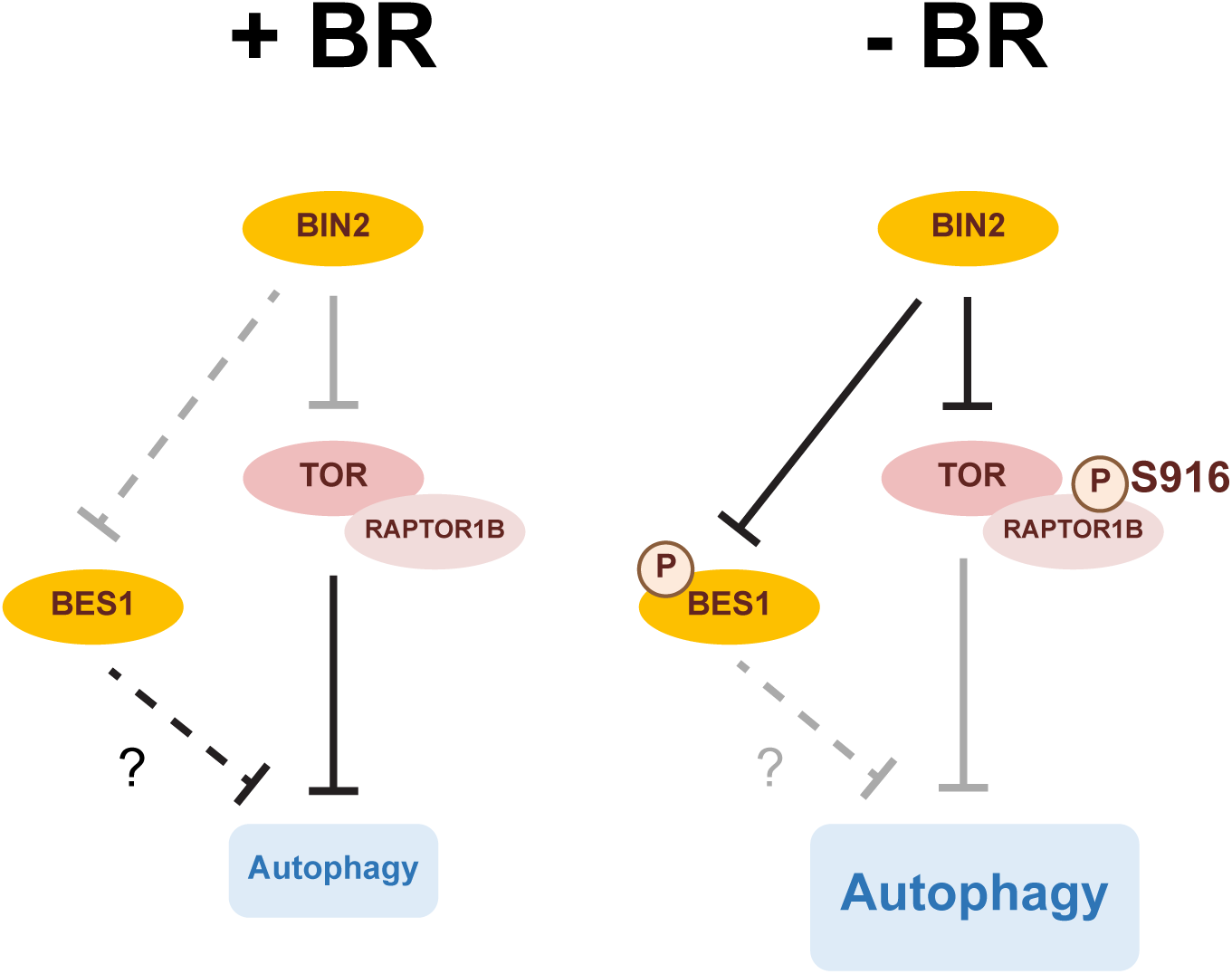
Proposed model for autophagy regulation by BIN2-mediated phosphorylation of RAPTOR1B. In the presence of BRs, enhanced BR signaling prevents phosphorylation of RAPTOR1B and BES1 by BIN2, enhancing TOR activity, in turn reducing the autophagy level. In the absence of BRs, BIN2 phosphorylates both RAPTOR1B and BES1. The repression of BR signaling leads to activated autophagy.

S6K phosphorylation at Thr449 has been considered a reliable downstream readout for assessing TOR activity in Arabidopsis ^29^. Considering that several feedback loops between mTOR, GSKs, and S6K proteins have been described in mammalian cells, detecting a single target may not capture the true picture of TOR pathway activity due to these complex signaling interactions. As a key metabolic hub, TORC controls multiple downstream substrates in response to environmental cues. In mammals, mTORC1 phosphorylates transcription factor EB (TFEB), a major regulator of lysosomal biogenesis and autophagy, via a substrate-specific mechanism that is mediated by Rag GTPases ^76^. However, whether plant TORC responds to diverse stimuli by selectively phosphorylating specific substrates remains unknown. Our findings suggest that BIN2-mediated phosphorylation of RAPTOR1B may selectively fine-tune TOR activity toward substrates that regulate autophagy but have less effect on growth, which provides evidence that selective activity of TOR towards different substrates exists in plants. Further analysis of BR-mediated regulation of autophagy is needed to understand the roles of BIN2 in TOR-dependent/-independent autophagy regulation induced by different stresses, as well as to identify other BIN2 or TOR direct substrates as autophagy effectors.

Our results reveal evidence of phosphorylation of RAPTOR1B by a GSK3 family member, which contributes to autophagy regulation in plants. Considering that both TOR and GSK3 are critical targets of many human disease therapies, the autophagy regulatory mechanism we revealed here in Arabidopsis may be conserved in other plants and mammals that include this conserved serine. These discoveries may contribute to advancing knowledge of how plants regulate growth and stress responses independently, which is important for developing strategies to optimize plant adaptation to environmental changes.

## Materials and methods

### Plant materials and growth conditions

Mutants and transgenic lines used in this study include: *raptor1b* (salk_078159) ^21^, *GFP-ATG8e* ^77^, *TOROE* (G548) ^13^, *bri1-301* ^43, 78^, *bin2-1D* ^37^, and *bes1-D* ^79^. The *bes1-D bin2-D*, *bes1-D raptor1b,* and *TOROE bin2-D* double mutants were generated by crossing the corresponding single mutants. Homozygous lines were selected by PCR genotyping.

Arabidopsis seeds of WT (Col-0) or the above genotypes were sterilized with 33% (v/v) bleach and 0.1% (v/v) Triton X-100 for 20 min, followed by 5 washes of 5 minutes each with sterile water. Sterilized seeds were stored at 4°C in darkness for at least 2 days to allow stratification before plating on solid ½ MS medium (2.22 g/L Murashige-Skoog vitamin and salt mixture (MS; Caisson Laboratory, MSP01), 1% (w/v) sucrose, 0.6% (w/v) Phytoblend agar (Caisson Laboratory, PTP01), 2.4 mM 2-morphinolino-ethanesulfonic acid (MES; Sigma-Aldrich, M3671), pH 5.7). Seedlings were grown in long-day conditions (16 h light) at 22°C for 7 days. Plants for transient expression in leaf protoplasts were grown in soil in a humidity-controlled growth chamber with 50% humidity at 20-23°C under long-day conditions for 4 to 6 weeks. For sucrose starvation, 7-day-old seedlings grown on solid ½ MS medium were transferred to solid ½ MS medium lacking sucrose and kept in darkness for an additional 3 days.

### BR and BRZ response assays

Sterilized Arabidopsis seeds were germinated and grown on solid ½ MS medium with DMSO, BL or AZD8055 at the indicated concentrations in the light for 7 days. For BRZ response assays, sterilized seeds were plated on solid ½ MS medium with DMSO or the indicated concentrations of BRZ. Plates were exposed to light for 6-8 h and then kept in darkness for 7 days ^8^. Plates were imaged using the Epson Perfection V600 Flatbed Photo scanner and hypocotyl lengths were measured using ImageJ ^80^.

### Gene expression analysis

Previously reported RNA-seq data for 4-week-old WT plants treated with or without BL or AZD8055 were used to analyze the effect of BL on TOR-regulated genes ^58, 81^. TOR- and BL-regulated gene lists were compared using Venny (http://bioinfogp.cnb.csic.es/tools/venny/index.html). Significance of observed gene overlaps was assessed via a hypergeometric test. Clustering analysis and heatmap construction was performed using the ‘aheatmap’ function of the NMF package in R (https://cran.r-project.org/web/packages/NMF/index.html) using log2 reads per million mapped reads (RPM) values.

### Western blot

Protein extracts from Arabidopsis seedlings treated with BL or AZD8055 were prepared in 2x Laemmli sample buffer with β-mercaptoethanol, separated by SDS-PAGE, and transferred to nitrocellulose membrane. Immunoblotting was performed using polyclonal rabbit antibody against BES1 ^55^, HERK1 ^82^, S6K1/2 (Agrisera, AS12 1855), P-T449-S6K (Agrisera, AS13 2664), monoclonal anti-FLAG (Sigma-Aldrich, F1804) or monoclonal anti-GFP (Sigma-Aldrich, MAB3836) and then with horseradish peroxidase–conjugated Goat Anti-Rabbit IgG antibodies (1:15000 dilution, Bio-Rad, 1706515) or Goat Anti-Mouse antibody (1:15000 dilution; Invitrogen, A10677). Signals were detected using chemiluminescence.

### Autophagy detection by fluorescence microscopy

*GFP-ATG8e* transgenic seedlings were observed and photographed using a Zeiss Axio Imager.A2 upright microscope (Zeiss, Oberkochen, Germany) equipped with Zeiss Axiocam BW/color digital cameras and a GFP-specific filter at the Iowa State University Microscopy and Nanoimaging Facility. Cells within the root elongation zone were photographed and the number of autophagosomes in each image was counted and averaged from at least 10 images per sample ^83^. For monodansylcadaverine (MDC) staining, Arabidopsis seedling roots were stained with MDC (Sigma-Aldrich, 30432) as described previously ^63^. MDC-stained seedlings were observed and imaged with the same fluorescence microscopy system using a DAPI-specific filter.

### Transient expression in protoplasts

*GFP-ATG8e* or *mCherry-ATG8e* ^84^ was transiently expressed alone in Arabidopsis leaf protoplasts as previously described ^85, 86^, or with YFP-BIN2-D ^8^ or YFP-BES1-D constructs ^8^. 25-30 μg of each plasmid DNA was introduced into protoplasts using 40% (w/v) polyethylene glycol. Protoplasts were washed and incubated in W5 solution (154 mM NaCl, 125 mM CaCl_2_, 5 mM KCl, 2 mM MES, pH 5.7) without sucrose for starvation stress, or with 0.5% (w/v) sucrose as control at room temperature in darkness for 2 days. Protoplasts were observed by Nikon Eclipse E200 fluorescence microscopy (Nikon Instruments Inc., New York, U.S.A.) using a GFP-specific filter for YFP and GFP, and an RFP-specific filter for mCherry. Protoplasts with 3 or more visible autophagosomes were counted as active for autophagy ^87^. A total of 100 protoplasts were observed per genotype for each condition, and the percentage of protoplasts with induced autophagy was calculated and averaged from 3 independent experimental replicates.

For ATG13a phosphorylation assays, ATG13a-FLAG was transiently expressed in Arabidopsis leaf protoplasts from each genotype using the same method described above for 18 hours, and 1 μM concanamycin A (ConA) (Sigma-Aldrich, C9705) or DMSO as control was added for 12 hours before collecting samples for immunoblotting analysis.

### Generation of constructs and transgenic plants

Constructs were generated via restriction enzyme digestion or Gateway technology and confirmed by DNA sequencing. The *RAPTOR1B* coding region was divided into 3 fragments for production in *E. coli*, and each fragment was amplified from Col-0 cDNA with primers including restriction sites (Table S1). Fragments were digested with EcoRI and SalI. Digested fragments were ligated into the pET-MBP-H vector ^8^, introducing a MBP tag at the N-terminus of each. YFP-BES1-D was generated via Gateway recombination with the pDH51-YFP destination vector. RAPTOR1B full-length cDNA was cloned into pENTR4 and used for generating site-mutated constructs with the Q5 site-directed mutagenesis kit (New England Biolabs, E0554S). *RAPTOR1B* full-length and site-mutated cDNA were subcloned into binary vector pGWB611 by Gateway LR recombination reaction (Invitrogen, 11791020). Primers used in this study are listed in Table S1. Plasmids were transferred into *Agrobacterium tumefaciens* GV3101 and used to transform plants by the floral-dip method ^88^. Transgenic plants were screened for antibiotic resistance and confirmed by RT-PCR. For ATG13a-FLAG, ATG13a cDNA was amplified by the same method described above, then digested with BamHI and SalI. The digested fragment was ligated into the pPZP211 vector ^45^, introducing a FLAG tag at the C-terminus.

### RNA extraction and quantitative real-time PCR

Total RNA was extracted from transgenic seedlings carrying 35S promoter-driven wild-type *RAPTOR1B/ raptor1b* or *RAPTOR1B^S916A^/raptor1b* and control seedlings using a Qiagen RNeasy kit (Qiagen, 74104) according to the manufacturer’s instructions. The first strand cDNA was synthesized by Superscript II reverse transcriptase (Invitrogen, 18080044). For quantitative real-time PCR assays, iQ SYBR Green Supermix (Bio-Rad, 1708880) was used for reaction set up. Applied Biosystems StepOnePlus™ was used to detect amplification levels (initial step at 95°C for 5 min, followed by 40 cycles of 15 s at 95°C, 1 min at 60°C and 1 min at 72°C). Results were normalized to the reference gene Actin2 using the ΔΔCt method. Each experiment was repeated using three independent biological replicates. Primers used in this study are listed in Table S1.

### *in vitro* protein-protein interaction and phosphorylation assay

Recombinant proteins were produced in *E. coli* strain BL21, and GST-pull down experiments were carried out as described previously ^45^. MBP or each MBP-tagged RAPTOR1B fragment was incubated with GST-BIN2 beads in 200 µl GST-pulldown buffer (50 mM Tris-HCl pH 7.5, 150 mM NaCl, 1 mM EDTA, 1 mM DTT, and 0.5% (v/v) NP-40) including 1 mg/mL BSA at 4 °C for 2 hours on a tube rotator. GST beads were washed in GST-pulldown buffer 4 times and then eluted in 2X SDS sample buffer. Each MBP-tagged RAPTOR1B fragment was detected with anti-MBP antibodies (NEB, E8032S). For the *in vitro* kinase assay, each MBP-tagged RAPTOR1B fragment, or MBP alone as control, was incubated with MBP-BIN2 kinase in 20 μL of kinase buffer (20 mM Tris, pH 7.5, 100 mM NaCl, 12 mM MgCl_2_ and 10 µCi ^32^P-ATP) and *in vitro* kinase assays carried out as described previously^45^.

### Statistical analysis

The significance of differences between means was analyzed by Fisher’s exact test or by Student’s *t*-test with *p* < 0.05 denoted *, and *p* < 0.01 denoted **, and *p* < 0.001 denoted ***, or with different letters to indicate statistically significant differences (*p* < 0.05).

## Author contributions

YP, TN, YY, JW and DCB conceived the project; YP and TN performed initial investigations; DCB, YY, JW acquired funding; CYL performed experiments with the following exceptions: YP performed the experiments in Figure 1C-F, Figure S1, Figure 2A and E, Figure S3A-B; TN conducted the RNA-seq data analysis in Figure 1A-B; CM and TN performed the phosphoproteomics, MS data analysis and statistical analysis of the phosphoproteome for identification of RAPTOR1B phosphorylation sites in Figure 4A; YP and TN wrote a first draft of the manuscript; CYL and DCB wrote the manuscript; all authors provided intellectual insight and revised and approved the manuscript.

## Acknowledgements

We thank Dr. Maureen Hanson for providing *raptor1b* mutant seeds and Margaret Carter for assistance with confocal microscopy.

## References

1. Liao CY, Bassham DC. Combating stress: the interplay between hormone signaling and autophagy in plants. J Exp Bot 2020; 71:1723–33.

2. Marshall RS, Vierstra RD. Autophagy: the master of bulk and selective recycling. Annu Rev Plant Biol 2018; 69:173–208.

3. Zhao YG, Zhang H. Formation and maturation of autophagosomes in higher eukaryotes: a social network. Curr Opin Cell Biol 2018; 53:29–36.

4. Tsukada M, Ohsumi Y. Isolation and characterization of autophagy-defective mutants of Saccharomyces cerevisiae. FEBS Lett 1993; 333:169–74.

5. Farre JC, Subramani S. Mechanistic insights into selective autophagy pathways: lessons from yeast. Nat Rev Mol Cell Biol 2016; 17:537–52.

6. Birgisdottir AB, Lamark T, Johansen T. The LIR motif - crucial for selective autophagy. J Cell Sci 2013; 126:3237–47.

7. Xie Q, Tzfadia O, Levy M, Weithorn E, Peled-Zehavi H, Van Parys T, et al. hfAIM: A reliable bioinformatics approach for in silico genome-wide identification of autophagy-associated Atg8-interacting motifs in various organisms. Autophagy 2016; 12:876–87.

8. Nolan TM, Brennan B, Yang M, Chen J, Zhang M, Li Z, et al. Selective autophagy of BES1 mediated by DSK2 balances plant growth and survival. Dev Cell 2017; 41:33–46 e7.

9. Pu Y, Luo X, Bassham DC. TOR-dependent and -independent pathways regulate autophagy in Arabidopsis thaliana. Front Plant Sci 2017; 8:1204.

10. Laureano-Marin AM, Aroca A, Perez-Perez ME, Yruela I, Jurado-Flores A, Moreno I, et al. Abscisic Acid-Triggered Persulfidation of the Cys Protease ATG4 Mediates Regulation of Autophagy by Sulfide. Plant Cell 2020; 32:3902–20.

11. Acheampong AK, Shanks C, Cheng CY, Schaller GE, Dagdas Y, Kieber JJ. EXO70D isoforms mediate selective autophagic degradation of type-A ARR proteins to regulate cytokinin sensitivity. Proc Natl Acad Sci U S A 2020; 117:27034–43.

12. Tarnowski L, Rodriguez MC, Brzywczy J, Piecho-Kabacik M, Krckova Z, Martinec J, et al. A selective autophagy cargo receptor NBR1 modulates abscisic acid signalling in Arabidopsis thaliana. Sci Rep 2020; 10:7778.

13. Deprost D, Yao L, Sormani R, Moreau M, Leterreux G, Nicolai M, et al. The Arabidopsis TOR kinase links plant growth, yield, stress resistance and mRNA translation. EMBO Rep 2007; 8:864–70.

14. Liu Y, Bassham DC. TOR is a negative regulator of autophagy in Arabidopsis thaliana. PLoS One 2010; 5:e11883.

15. Noda T, Ohsumi Y. Tor, a phosphatidylinositol kinase homologue, controls autophagy in yeast. J Biol Chem 1998; 273:3963–6.

16. Pattingre S, Espert L, Biard-Piechaczyk M, Codogno P. Regulation of macroautophagy by mTOR and Beclin 1 complexes. Biochimie 2008; 90:313–23.

17. Dobrenel T, Caldana C, Hanson J, Robaglia C, Vincentz M, Veit B, et al. TOR Signaling and Nutrient Sensing. Annu Rev Plant Biol 2016; 67:261–85.

18. Wu Y, Shi L, Li L, Fu L, Liu Y, Xiong Y, et al. Integration of nutrient, energy, light and hormone signalling via TOR in plants. J Exp Bot 2019; 70:2227–38.

19. Yang H, Rudge DG, Koos JD, Vaidialingam B, Yang HJ, Pavletich NP. mTOR kinase structure, mechanism and regulation. Nature 2013; 497:217–23.

20. Hara K, Maruki Y, Long X, Yoshino K, Oshiro N, Hidayat S, et al. Raptor, a binding partner of target of rapamycin (TOR), mediates TOR action. Cell 2002; 110:177–89.

21. Anderson GH, Veit B, Hanson MR. The Arabidopsis AtRaptor genes are essential for post-embryonic plant growth. BMC Biol 2005; 3:12.

22. Deprost D, Truong HN, Robaglia C, Meyer C. An Arabidopsis homolog of RAPTOR/KOG1 is essential for early embryo development. Biochem Biophys Res Commun 2005; 326:844–50.

23. Menand B, Desnos T, Nussaume L, Berger F, Bouchez D, Meyer C, et al. Expression and Disruption of the Arabidopsis TOR (Target of Rapamycin) Gene. Proc Natl Acad Sci U S A 2002; 99:6422–7.

24. Ren M, Qiu S, Venglat P, Xiang D, Feng L, Selvaraj G, et al. Target of rapamycin regulates development and ribosomal RNA expression through kinase domain in Arabidopsis. Plant Physiol 2011; 155:1367.

25. Anderson GH, Hanson MR. The Arabidopsis Mei2 homologue AML1 binds AtRaptor1B, the plant homologue of a major regulator of eukaryotic cell growth. BMC Plant Biol 2005; 5:2.

26. Moreau M, Azzopardi M, Clement G, Dobrenel T, Marchive C, Renne C, et al. Mutations in the Arabidopsis homolog of LST8/GbetaL, a partner of the target of rapamycin kinase, impair plant growth, flowering, and metabolic adaptation to long days. Plant Cell 2012; 24:463–81.

27. Salem MA, Li Y, Bajdzienko K, Fisahn J, Watanabe M, Hoefgen R, et al. RAPTOR controls developmental growth transitions by altering the hormonal and metabolic balance. Plant Physiol 2018; 177:565–93.

28. Mahfouz MM, Kim S, Delauney AJ, Verma DPS. Arabidopsis TARGET OF RAPAMYCIN interacts with RAPTOR, which regulates the activity of S6 kinase in response to osmotic stress signals. Plant Cell 2006; 18:477–90.

29. Xiong Y, Sheen J. Rapamycin and glucose-target of rapamycin (TOR) protein signaling in plants. The Journal of biological chemistry 2012; 287:2836.

30. Van Leene J, Han C, Gadeyne A, Eeckhout D, Matthijs C, Cannoot B, et al. Capturing the phosphorylation and protein interaction landscape of the plant TOR kinase. Nat Plants 2019; 5:316–27.

31. Suttangkakul A, Li F, Chung T, Vierstra RD. The ATG1/ATG13 protein kinase complex is both a regulator and a target of autophagic recycling in Arabidopsis. Plant Cell 2011; 23:3761–79.

32. Xiong FJ, Zhang R, Meng ZG, Deng KX, Que YM, Zhuo FP, et al. Brassinosteriod Insensitive 2 (BIN2) acts as a downstream effector of the Target of Rapamycin (TOR) signaling pathway to regulate photoautotrophic growth in Arabidopsis. New Phytol 2017; 213:233–49.

33. Zhang Z, Zhu JY, Roh J, Marchive C, Kim SK, Meyer C, et al. TOR signaling promotes accumulation of BZR1 to balance growth with carbon availability in Arabidopsis. Curr Biol 2016; 26:1854–60.

34. Nolan TM, Vukasinovic N, Liu D, Russinova E, Yin Y. Brassinosteroids: Multidimensional Regulators of Plant Growth, Development, and Stress Responses. Plant Cell 2020; 32:295–318.

35. Clouse SD, Langford M, McMorris TC. A brassinosteroid-insensitive mutant in Arabidopsis thaliana exhibits multiple defects in growth and development. Plant Physiol 1996; 111:671–8.

36. Li J, Chory J. A Putative Leucine-Rich Repeat Receptor Kinase Involved in Brassinosteroid Signal Transduction. Cell 1997; 90:929–38.

37. Li J, Nam KH, Vafeados D, Chory J. BIN2, a new Brassinosteroid-insensitive locus in Arabidopsis. Plant Physiol 2001; 127:14.

38. Li J, Wen J, Lease KA, Doke JT, Tax FE, Walker JC. BAK1, an Arabidopsis LRR receptor-like protein kinase, interacts with BRI1 and modulates brassinosteroid signaling. Cell 2002; 110:213–22.

39. Nam KH, Li J. BRI1/BAK1, a receptor kinase pair mediating brassinosteroid signaling. Cell 2002; 110:203–12.

40. Clouse SD. Brassinosteroid signal transduction: from receptor kinase activation to transcriptional networks regulating plant development. Plant Cell 2011; 23:1219–30.

41. Kim TW, Wang ZY. Brassinosteroid signal transduction from receptor kinases to transcription factors. Annu Rev Plant Biol 2010; 61:681–704.

42. Mao J, Li J. Regulation of Three Key Kinases of Brassinosteroid Signaling Pathway. Int J Mol Sci 2020; 21.

43. Li J, Nam KH. Regulation of Brassinosteroid Signaling by a GSK3/SHAGGY-like Kinase. Science 2002; 295:1299–301.

44. Wang ZY, Nakano T, Gendron J, He JX, Chen M, Vafeados D, et al. Nuclear-localized BZR1 mediates brassinosteroid-induced growth and feedback suppression of brassinosteroid biosynthesis. Dev Cell 2002; 2:505–13.

45. Yin Y, Wang Z-Y, Mora-Garcia S, Li J, Yoshida S, Asami T, et al. BES1 Accumulates in the Nucleus in Response to Brassinosteroids to Regulate Gene Expression and Promote Stem Elongation. Cell 2002; 109:181–91.

46. Zhao J, Peng P, Schmitz RJ, Decker AD, Tax FE, Li J. Two putative BIN2 substrates are nuclear components of brassinosteroid signaling. Plant Physiol 2002; 130:1221–9.

47. Clark NM, Nolan TM, Wang P, Song G, Montes C, Valentine CT, et al. Integrated omics networks reveal the temporal signaling events of brassinosteroid response in Arabidopsis. Nat Commun 2021; 12:5858.

48. He JX, Gendron JM, Yang YL, Li JM, Wang ZY. The GSK3-like kinase BIN2 phosphorylates and destabilizes BZR1, a positive regulator of the brassinosteroid signaling pathway in Arabidopsis. Proc Natl Acad Sci U S A 2002; 99:10185–90.

49. Vert G, Chory J. Downstream nuclear events in brassinosteroid signalling. Nature 2006; 441:96–100.

50. Gampala SS, Kim TW, He JX, Tang W, Deng Z, Bai MY, et al. An essential role for 14-3-3 proteins in brassinosteroid signal transduction in Arabidopsis. Dev Cell 2007; 13:177–89.

51. Tang W, Yuan M, Wang R, Yang Y, Wang C, Oses-Prieto JA, et al. PP2A activates brassinosteroid-responsive gene expression and plant growth by dephosphorylating BZR1. Nat Cell Biol 2011; 13:124–31.

52. Sun Y, Fan XY, Cao DM, Tang W, He K, Zhu JY, et al. Integration of brassinosteroid signal transduction with the transcription network for plant growth regulation in Arabidopsis. Dev Cell 2010; 19:765–77.

53. Ye HX, Liu SZ, Tang BY, Chen JN, Xie ZL, Nolan TM, et al. RD26 mediates crosstalk between drought and brassinosteroid signalling pathways. Nature Communications 2017; 8:14573.

54. Yin Y, Vafeados D, Tao Y, Yoshida S, Asami T, Chory J. A new class of transcription factors mediates brassinosteroid-regulated gene expression in Arabidopsis. Cell 2005; 120:249–59.

55. Yu X, Li L, Zola J, Aluru M, Ye H, Foudree A, et al. A brassinosteroid transcriptional network revealed by genome-wide identification of BESI target genes in Arabidopsis thaliana. Plant J 2011; 65:634–46.

56. He JX, Gendron JM, Sun Y, Gampala SSL, Gendron N, Sun CQ, et al. BZR1 is a transcriptional repressor with dual roles in brassinosteroid homeostasis and growth responses. Science 2005; 307:1634–8.

57. Guo H, Li L, Aluru M, Aluru S, Yin Y. Mechanisms and networks for brassinosteroid regulated gene expression. Curr Opin Plant Biol 2013; 16:545–53.

58. Dong P, Xiong FJ, Que YM, Wang K, Yu LH, Li ZG, et al. Expression profiling and functional analysis reveals that TOR is a key player in regulating photosynthesis and phytohormone signaling pathways in Arabidopsis. Frontiers in Plant Science 2015; 6:677.

59. Wang X, Chen J, Xie Z, Liu S, Nolan T, Ye H, et al. Histone lysine methyltransferase SDG8 is involved in brassinosteroid-regulated gene expression in Arabidopsis thaliana. Mol Plant 2014; 7:1303–15.

60. Xiong Y, McCormack M, Li L, Hall Q, Xiang C, Sheen J. Glucose-TOR signalling reprograms the transcriptome and activates meristems. Nature 2013; 496:181.

61. Asami T, Min YK, Nagata N, Yamagishi K, Takatsuto S, Fujioka S, et al. Characterization of brassinazole, a triazole-type brassinosteroid biosynthesis inhibitor. Plant Physiol 2000; 123:93–100.

62. Mizushima N, Yoshimori T, Levine B. Methods in mammalian autophagy research. Cell 2010; 140:313–26.

63. Contento AL, Xiong Y, Bassham DC. Visualization of autophagy in Arabidopsis using the fluorescent dye monodansylcadaverine and a GFP-AtATG8e fusion protein. The Plant journal : for cell and molecular biology 2005; 42:598.

64. Rodriguez E, Chevalier J, Olsen J, Ansbol J, Kapousidou V, Zuo Z, et al. Autophagy mediates temporary reprogramming and dedifferentiation in plant somatic cells. EMBO J 2020; 39:e103315.

65. Youn JH, Kim TW. Functional insights of plant GSK3-like kinases: multi-taskers in diverse cellular signal transduction pathways. Mol Plant 2015; 8:552–65.

66. Montes C, Liao C-Y, Nolan TM, Song G, Clark NM, Guo H, et al. Interplay between brassinosteroids and TORC signaling in Arabidopsis revealed by integrated multi-dimensional analysis. bioRxiv 2021:2021.02.12.431003.

67. Jayaraman D, Richards AL, Westphall MS, Coon JJ, Ane JM. Identification of the phosphorylation targets of symbiotic receptor-like kinases using a high-throughput multiplexed assay for kinase specificity. Plant J 2017; 90:1196–207.

68. Brumbaugh J, Russell JD, Yu P, Westphall MS, Coon JJ, Thomson JA. NANOG is multiply phosphorylated and directly modified by ERK2 and CDK1 in vitro. Stem Cell Reports 2014; 2:18–25.

69. Aylett CH, Sauer E, Imseng S, Boehringer D, Hall MN, Ban N, et al. Architecture of human mTOR complex 1. Science 2016; 351:48–52.

70. Veit B, Anderson Garrett H, Hanson Maureen R. The Arabidopsis AtRaptor genes are essential for post-embryonic plant growth. BMC Biol 2005; 3:12.

71. Henriques R, Bögre L, Horváth B, Magyar Z. Balancing act: matching growth with environment by the TOR signalling pathway. J Exp Bot 2014; 65:2691–701.

72. Chaiwanon J, Wang W, Zhu JY, Oh E, Wang ZY. Information Integration and Communication in Plant Growth Regulation. Cell 2016; 164:1257–68.

73. Doble BW, Woodgett JR. GSK-3: tricks of the trade for a multi-tasking kinase. J Cell Sci 2003; 116:1175–86.

74. Yan Z, Zhao J, Peng P, Chihara RK, Li J. BIN2 functions redundantly with other Arabidopsis GSK3-like kinases to regulate brassinosteroid signaling. Plant Physiol 2009; 150:710–21.

75. Stretton C, Hoffmann TM, Munson MJ, Prescott A, Taylor PM, Ganley IG, et al. GSK3-mediated raptor phosphorylation supports amino-acid-dependent mTORC1-directed signalling. Biochem J 2015; 470:207–21.

76. Napolitano G, Di Malta C, Esposito A, de Araujo MEG, Pece S, Bertalot G, et al. A substrate-specific mTORC1 pathway underlies Birt-Hogg-Dube syndrome. Nature 2020; 585:597–602.

77. Xiong Y, Contento AL, Nguyen PQ, Bassham DC. Degradation of Oxidized Proteins by Autophagy during Oxidative Stress in Arabidopsis. Plant Physiol 2007; 143:291–9.

78. Xu W, Huang J, Li B, Li J, Wang Y. Is kinase activity essential for biological functions of BRI1? Cell Res 2008; 18:472–8.

79. Vilarrasa-Blasi J, Gonzalez-Garcia MP, Frigola D, Fabregas N, Alexiou KG, Lopez-Bigas N, et al. Regulation of plant stem cell quiescence by a brassinosteroid signaling module. Dev Cell 2014; 30:36–47.

80. Schneider CA, Rasband WS, Eliceiri KW. NIH Image to ImageJ: 25 years of image analysis. Nat Methods 2012; 9:671–5.

81. Wang W, Bai MY, Wang ZY. The brassinosteroid signaling network-a paradigm of signal integration. Curr Opin Plant Biol 2014; 21:147–53.

82. Guo H, Ye H, Li L, Yin Y. A family of receptor-like kinases are regulated by BES1 and involved in plant growth in Arabidopsis thaliana. Plant Signal Behav 2009; 4:784–6.

83. Pu Y, Bassham DC. Detection of Autophagy in Plants by Fluorescence Microscopy. Methods Mol Biol 2016; 1450:161–72.

84. Bao Y, Song WM, Wang P, Yu X, Li B, Jiang C, et al. COST1 regulates autophagy to control plant drought tolerance. Proc Natl Acad Sci U S A 2020; 117:7482–93.

85. Yoo SD, Cho YH, Sheen J. Arabidopsis mesophyll protoplasts: a versatile cell system for transient gene expression analysis. Nat Protoc 2007; 2:1565–72.

86. Wu FH, Shen SC, Lee LY, Lee SH, Chan MT, Lin CS. Tape-Arabidopsis Sandwich - a simpler Arabidopsis protoplast isolation method. Plant Methods 2009; 5:16.

87. Yang X, Srivastava R, Howell SH, Bassham DC. Activation of autophagy by unfolded proteins during endoplasmic reticulum stress. Plant J 2016; 85:83–95.

88. Clough SJ, Bent AF. Floral dip: a simplified method for Agrobacterium-mediated transformation of Arabidopsis thaliana. Plant J 1998; 16:735–43.

