## Supplementary material for "Brassinosteroids modulate autophagy through phosphorylation of RAPTOR1B by the GSK3-like kinase BIN2 in Arabidopsis": Figure S1

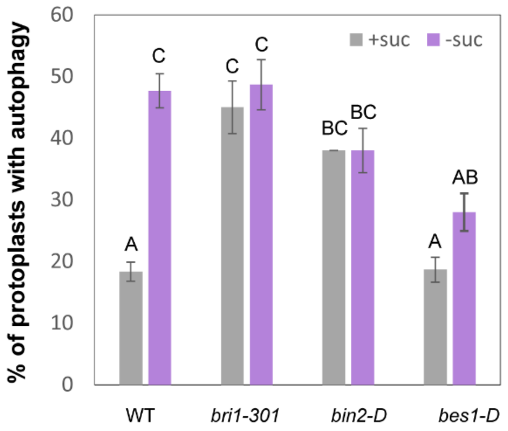


**Figure S1.** Defects in BR signaling lead to autophagy induction. Quantification of autophagy in protoplasts by calculating the percentage of protoplasts with 3 or more GFP-ATG8e-labeled autophagosomes in the indicated genotype in the presence or absence of sucrose. Data represent means ± SD from 3 biological replicates. Different letters indicate statistically significant differences ($p < 0.05$) using Student's *t*-test.

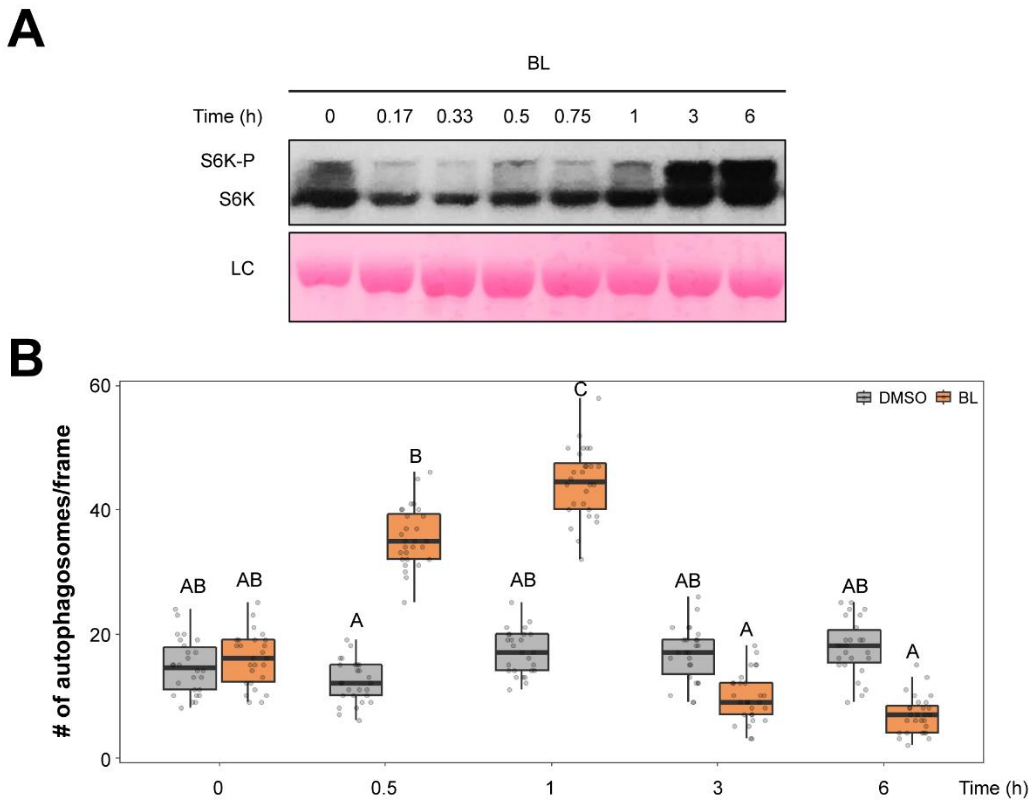


**Figure S2.** BL reactivates TOR, leading to autophagy inhibition after perception of stimuli. (**A**) Immunoblot of TOR-dependent S6K phosphorylation (S6K-P) in WT seedlings. Seedlings were grown on solid ½ MS medium for 7 days and transferred to liquid ½ MS medium with 100 nM BL for 0, 10, 20, 30, 45 minutes and 1, 3, 6 hours. Ponceau staining was used as a loading control. Experiments were repeated a minimum of 3 times with similar results. (**B**) Number of GFP-tagged autophagosomes per frame upon 0, 0.5, 1, 3, 6 hours of 100 nM BL treatment or DMSO as control. GFP-labeled autophagosomes were observed by fluorescence microscopy and photographed. The average number of autophagosomes was calculated from at least 20 images per genotype for each condition. Data represent means ± SD. Boxes show the corresponding mean of each replicate with round dots representing individual measurements from each replicate. Different letters indicate statistically significant differences ($p < 0.05$) using Student's *t*-test.

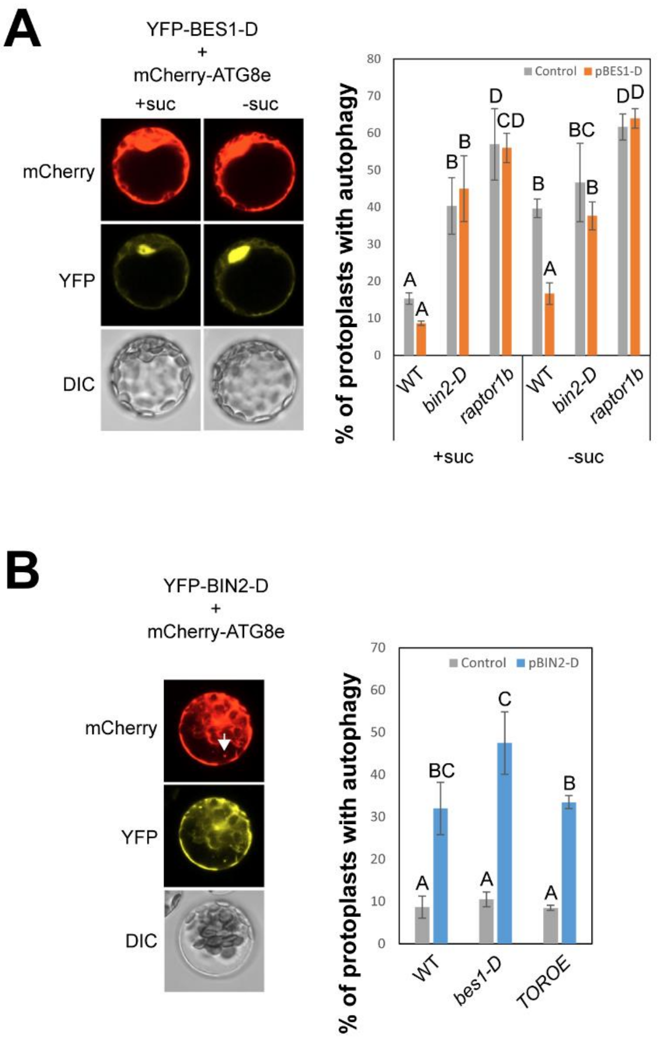


**Figure S3.** BIN2 activates autophagy in protoplasts and seedling roots. (**A**) Left panel, representative confocal images of protoplasts expressing mCherry-ATG8e and YFP-BES1-D. After transformation, leaf protoplasts were incubated in control (+suc) or starvation (-suc) conditions for 2 days. Right panel, quantification of autophagy in protoplasts by calculating the percentage of protoplasts with 3 or more mCherry-ATG8e-labeled autophagosomes when transiently overexpressing YFP-BES1-D (pBES1-D) in the indicated genotype for each condition. Data represent means ± SD from 3 biological replicates. (**B**) Left panel, representative confocal images of protoplasts expressing mCherry-ATG8e and YFP-BIN2-D.

Autophagic bodies are indicated by the arrows. Right panel, quantification of autophagy in protoplasts by calculating the percentage of protoplasts with 3 or more mCherry-ATG8e-labeled autophagosomes when transiently overexpressing YFP-BIN2-D (pBIN2-D) in the indicated genotype for each condition. Data represent means ± SD from 3 biological replicates.

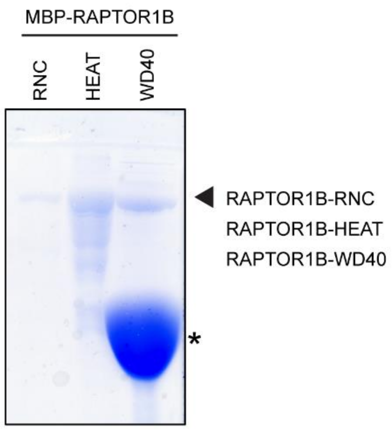


**Figure S4.** Coomassie blue staining of RAPTOR1B recombinant proteins. Asterisk indicates degraded RAPTOR1B-WD40 protein.

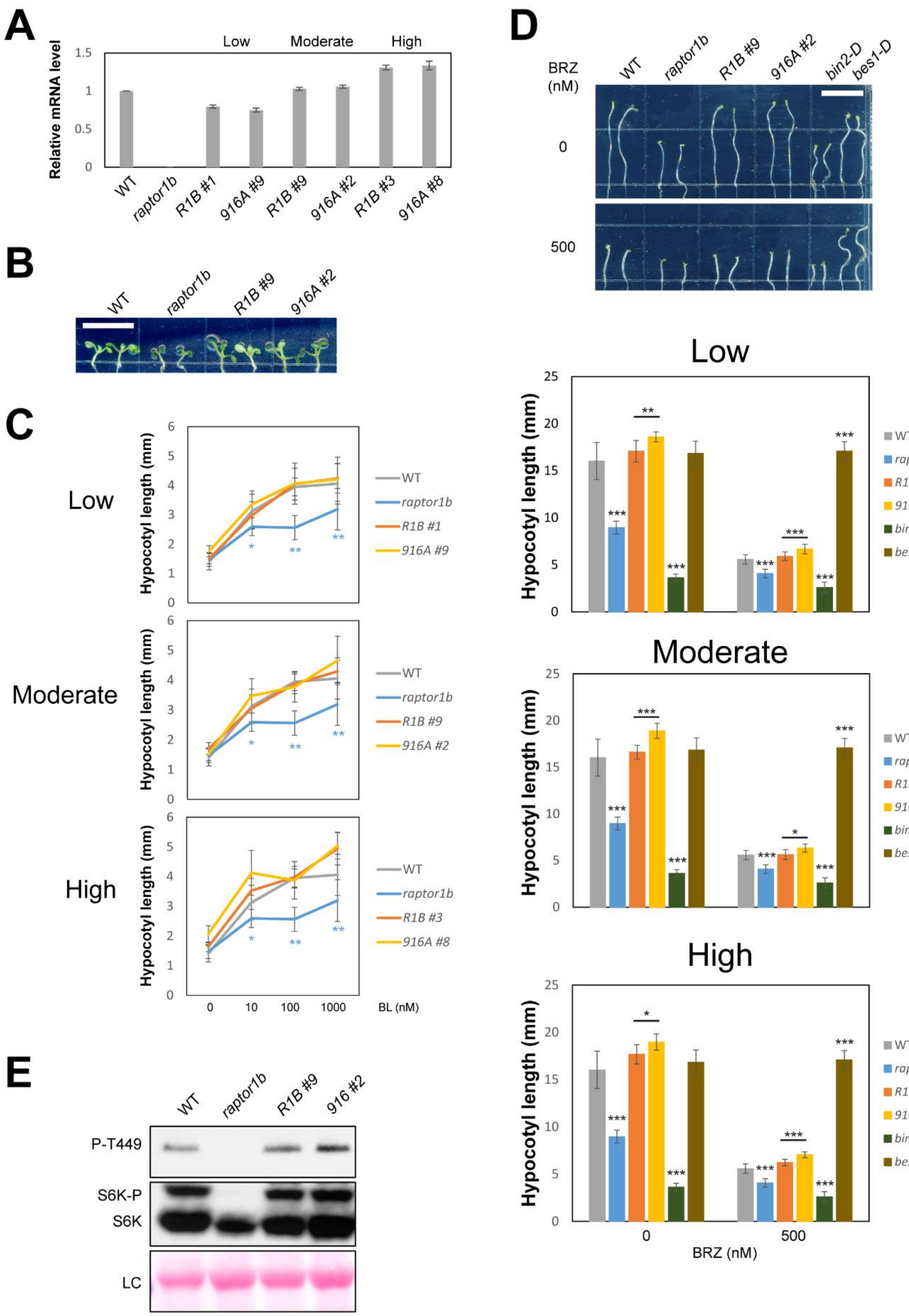

A
Low
Moderate
High
Relative mRNA level
1.5
1
0.5
0
WT
raptor1b
R1B #1
916A #9
R1B #9
916A #2
R1B #3
916A #8
B
WT
raptor1b
R1B #9
916A #2
C
Low
Moderate
High
Hypocotyl length (mm)
WT
raptor1b
R1B #1
916A #9
R1B #9
916A #2
R1B #3
916A #8
0
10
100
1000
BL (nM)
D
BRZ
(nM)
0
500
WT
raptor1b
R1B #9
916A #2
bin2-D
bes1-D
Low
Moderate
High
Hypocotyl length (mm)
WT
raptor1b
R1B #1
916A #9
bin2-D
bes1-D
R1B #9
916A #2
R1B #3
916A #8
BRZ (nM)
E
WT
raptor1b
R1B #9
916 #2
P-T449
S6K-P
S6K
LC

**Figure S5.** Response of RAPTOR1B seedlings in *raptor1b* background to BL and BRZ. (**A**) Expression of *RAPTOR1B* gene in *raptor1b* transgenic plants expressing WT RAPTOR1B or RAPTOR1B$^{S916A}$. Three independent lines of each genotype were used. Transgene expression levels of each line were measured using real-time PCR and normalized to the wild-type *RAPTOR1B* expression level in a wild-type background. Transgenic plants that have similar expression levels were grouped together (“Low”, “Moderate”, and “High”) for comparison. No difference in phenotype was seen between groups. (**B**) Representative 7-day-old hypocotyl length of each genotype. Seedlings were grown on ½ MS medium for 7 days. Scale bar = 10 mm. (**C**) Hypocotyl elongation of each genotype in response to BL. Seedlings of each genotype were grown on ½ MS medium with the indicated BL concentration for 7 days, and hypocotyl length measured. (**D**) Hypocotyl elongation of each genotype in response to dark and BRZ. Seedlings were grown on ½ MS medium with the indicated BRZ concentration in the dark for 7 days. Representative seedlings are shown in the upper panel. Scale bar = 10 mm. (**E**) Immunoblot of TOR-dependent S6K phosphorylation (P-T449 and S6K-P) in WT seedlings, *raptor1b*, and transgenic plants expressing *RAPTOR1B*$^{S916A}$ in a *raptor1b* background. Seedlings were grown on solid ½ MS medium for 7 days. Ponceau staining was used as a loading control. For panel (**C**) and (**D**), hypocotyl lengths of seedlings were measured and averaged from 15 independent seedlings and 3 independent replicates. Data represent means ± SD. Asterisks indicate statistically significant differences (***$p < 0.001$, **$p < 0.01$, *$p < 0.05$) using Student’s *t*-test compared with WT under the same conditions.

**A**

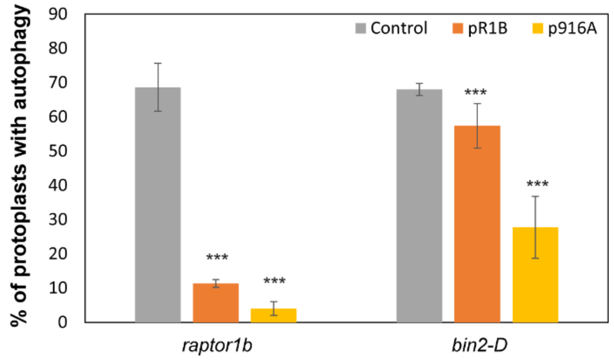
% of protoplasts with autophagy
Control
pR1B
p916A
***
raptor1b
bin2-D
0
10
20
30
40
50
60
70
80
90

**B**

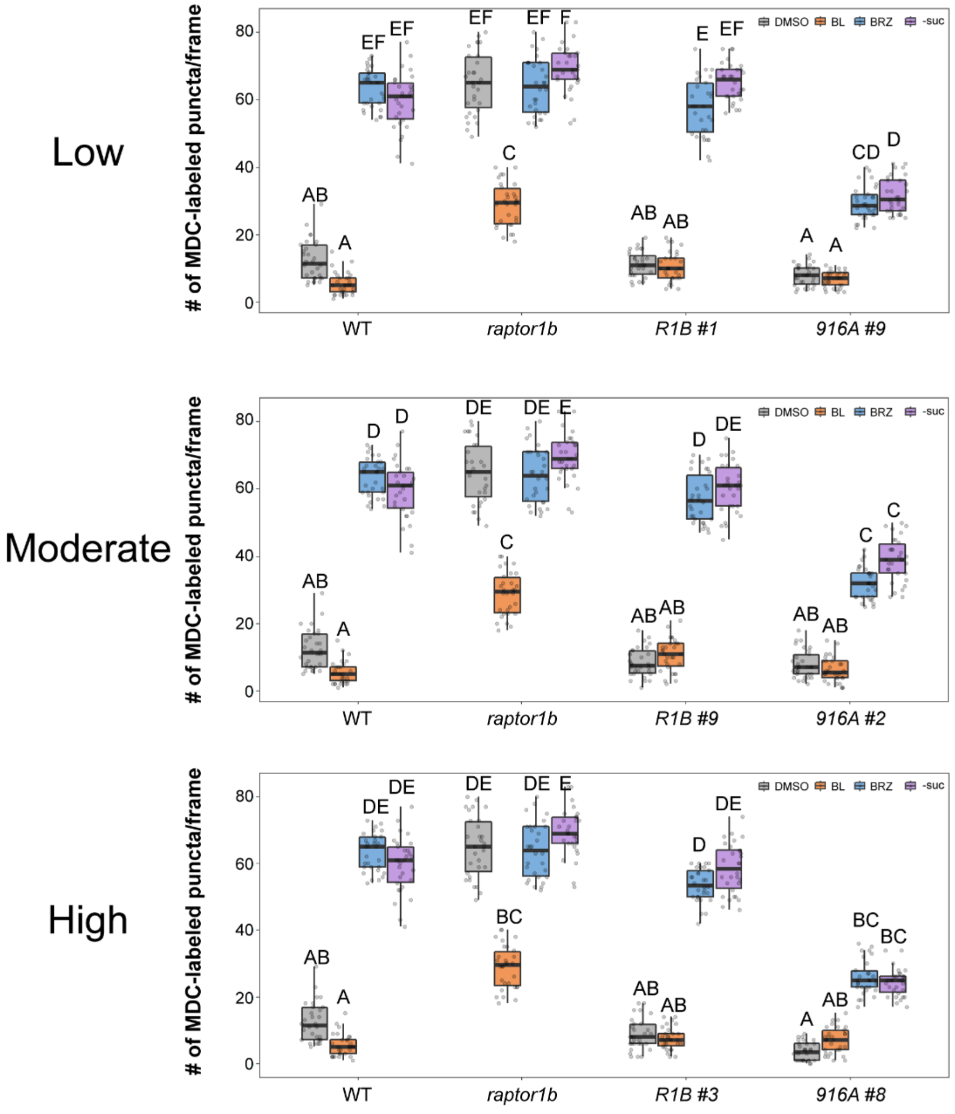
Low
Moderate
High
### of MDC-labeled puncta/frame
DMSO
BL
BRZ
-suc
WT
raptor1b
R1B #1
916A #9
R1B #9
916A #2
R1B #3
916A #8
A
AB
C
CD
D
DE
E
EF
F
BC
0
20
40
60
80

**Figure S6.** Autophagy response of RAPTOR1B$^{S916A}$ in different conditions. (**A**) Quantification of autophagy in protoplasts determined by calculating the percentage of protoplasts with 3 or more GFP-ATG8e-labeled autophagosomes when transiently overexpressing vector control, WT RAPTOR1B (R1B) or RAPTOR1B$^{S916A}$ (916A) in the indicated genotype. Asterisks indicate statistically significant differences (***$p < 0.001$) using Student's *t*-test compared with WT under the same conditions. (**B**) Number of MDC-stained puncta per frame in the indicated genotype for each condition. 7-day-old seedlings were transferred to solid ½ MS medium plus 100 nM BL, 1 μM BRZ, or DMSO as control, or under starvation conditions (-suc) for an additional 3 days. MDC-stained puncta were observed by fluorescence microscopy and photographed. The average number of puncta was calculated from at least 20 images per genotype for each condition. Different letters indicate statistically significant differences ($p < 0.05$) using Student's *t*-test. For all panels, data represent means ± SD. Boxes show the corresponding mean of each replicate with round dots representing individual measurements from each replicate.

**Table S1.** List of oligonucleotide primers.

| **Oligo name** | **Sequence** |
|---|---|
| R1B-EcoRIF | GCGgaattcATGGCATTAGGAGACTTAATGGTG |
| R1B-SalIR | CGGgtcgacTCATCTTGCTTGCGAGTTGTCGTG |
| WD40-EcoRIF | GCGgaattcTATTGCAAGTCTCGGACGGC |
| HEAT-EcoRIF | GCGgaattcTGGCTTGATCATGGATCTGA |
| HEAT-SalIR | CGGgtcgacTTAAAGCGGATCAGCCAATCCTG |
| RNC-SalIR | CGGgtcgacTTACAAGGCCAGATCCACAGCCC |
| R1BSalI-F | GCGgtcgacATGGCATTAGGAGACTTAATGGTG |
| R1BEcoRI-R | CGGgaattcTCATCTTGCTTGCGAGTTGTCGTGG |
| Q5-RAPTOR-S916A-F | TCCCCCGGTCgcgCCCCCTAGAAC |
| Q5-RAPTOR-S916A-R | GTCCTAAAACTTAAAGGCAG |
| Q5-RAPTOR-S916D-F | TCCCCCGGTCgaCCCCCCTAGA |
| Q5-RAPTOR-S916D-R | GTCCTAAAACTTAAAGGCAGATTACC |
| Q-R1B-F | ACTCACGGCCTTAGCTGTTC |
| Q-R1B-R | CTTGCTTGCGAGTTGTCGTG |
| ACTIN2F | GGAAGGATCTGTACGGTAAC |
| ACTIN2R | GGACCTGCCTCATCATACT |
| ATG13aBamHIF | CGCGGATCCATGGATTTTCCAGAGAATTTGCC |
| ATG13aSalIRns | ACGCGTCGACGTGGACGCGAGTTGGTCC |
